# Individual differences in prefrontal coding of visual features

**DOI:** 10.1101/2024.05.09.588948

**Authors:** Qi Lin, Hakwan Lau

## Abstract

Each of us perceives the world differently. What may underlie such individual differences in perception? Here, we characterize the lateral prefrontal cortex’s role in vision using computational models, with a specific focus on individual differences. Using a 7T fMRI dataset, we found that encoding models relating visual features extracted from a deep neural network to brain responses to natural images robustly predict responses in patches of LPFC. We then explored the representational structures and screened for images with high predicted responses in LPFC. We observed more substantial individual differences in the coding schemes of LPFC compared to visual regions. Computational modeling suggests that the amplified individual differences could result from the random projection between sensory and high-level regions underlying flexible working memory. Our study demonstrates the under-appreciated role of LPFC in visual processing and suggests that LPFC may underlie the idiosyncrasies in how different individuals experience the visual world.

## 1 Introduction

People experience the world differently from each other. Some of these individual differences could be attributed to high-level mental processes such as beliefs and reasoning (e.g., [1–3]). However, some of these idiosyncrasies in subjective experiences can also arise from the perceptual process itself. Here we investigate what may be the neural basis of these individual differences.

Situated at the apex of the cortical hierarchy, the prefrontal cortex (PFC) is disproportionally developed in primates [4]. The human PFC also exhibits higher variability across individuals in terms of both structural properties and functional connectivity architectures [5]. Moreover, the functional networks involving PFC are more varied across individuals and carry more information about individual differences in behavior (e.g., [6, 7]).

To date, PFC is most commonly associated with complex and high-level cognition, such as attention, language, and cognitive control[8–12]. The PFC is thought to underlie the impressive complexity and flexibility of human thoughts and intelligence. Empirical work has demonstrated that such flexibility is perhaps in part conferred by the prevalence of neurons demonstrating mixed selectivity in the PFC, in that these neurons respond to diverse combinations of experimental variables (e.g,, [13]; for reviews, see [14, 15]), in contrast to neurons in the early sensory cortex such as V1 that demonstrates relatively ‘pure’ or simple selectivity (e.g., cells responding to one orientation in a specific receptive field in [16]).

This distinction between the high prevalence of neurons with mixed selectivity in PFC compared to sensory cortex raises an intriguing question: despite its traditional association with high-level cognition due to the complexity of its coding scheme, could the PFC also play a significant role in basic perceptual functions? That is, could perception also benefit from circuits with high degree of mixed selectivity? Evidence from nonhuman primates has characterized the representation of visual information in PFC, especially the lateral PFC (LPFC), even when there is no cognitive task involved (for a review, see [17]). There are LPFC neurons showing selectivity towards a variety of visual features and categories during passive viewing (e.g., spatial location, shapes, and colors: [18]; faces: [19]; bodies, scenes, disparity and color: [20];category/image selectivity: [21, 22]). In addition to coding visual information in a way reminiscent of the visual cortex, LPFC are especially involved in visually challenging situations for nonhuman primates such as recognizing partially occluded objects [23] and are causally related to nonhuman primates’ successful recognition of images that are difficult for purely feedforward convolutional neural networks [24].

Despite the extensive evidence cited above from nonhuman primates for LPFC’s involvement in perception, to date, such detailed studies focusing on LPFC’s perceptual function in humans remain comparatively sparse. The content of subjective perception can be decoded from brain activities in LPFC when humans viewed ambiguous simple stimuli eliciting bistable perception (e.g., [25, 26]) or uniformity illusion [27]. PFC is causally related to such switching of subjective percepts [28]. In studies adopting a whole-brain approach with more complex stimuli such as scene images [29] and movies [30, 31], LPFC regions have often been found to be predictable based on semantic labels [30] or features extracted from deep neural networks [29, 31] but such results are often merely mentioned in passing without further investigation into how the coding in LPFC differs from the coding in sensory cortex. Admittedly, the relatively lack of attention given to these findings could be in part due to the relatively less robust nature of the relevant effects, as compared to those one could more readily observe within the visual cortex.

In this study, we aim to systematically characterize how the LPFC code visual stimuli, with a specific focus on individual differences. To this end, we built encoding models to relate visual features extracted from a deep neural network to predict brain activities in LPFC in an existing 7T human fMRI dataset [32]. We then contrasted the tuning profiles of LPFC for visual stimuli to those of the visual cortex and asked if the tuning profiles of LPFC demonstrate more individual variability. Lastly, we provide a simple mechanistic explanation of the difference between the tuning profiles of LPFC and those of visual cortex using computational modeling.

## 2 Results

Results presented here are based on data from the Natural Scenes Dataset (NSD; [32]). Each of the eight subjects viewed 9209 to 10,000 unique scene images and performed a continous recognition memory task while having their hemodynamic activities in the brain recorded in a 7T MRI scanner.

### 2.1 Encoding models robustly predict LPFC activities

To adequately characterize LPFC’s responses to naturalistic scene images and thus identify LPFC regions that show sensitivity to visual features, we reason that we need a feature space that is rich and expressive enough to describe the complexity of the images used in NSD and capture the multifaceted differences between these images. To this end, we turn to deep neural networks, inspired by previous works demonstrating the success of using features learned by deep neural networks in predicting activities in visual cortex (for reviews, see [33–35]). More specifically, to build encoding models of LPFC within each subject (see Figure 1A for a schematic of the prediction pipeline), we first extracted activations (vector length: 512) from the image encoder of a CLIP (Contranstive Language-Image Pretraining) network (CLIP-img, from ViT-B/32 backbone; [36]). The CLIP network was jointly trained with pairs of images and captions to minimize the distance between image and textual embeddings corresponding to the same pairs. In preliminary work, we also explored another network with a different architecture, ResNet-50[37] and saw qualitatively similar results. We decided to focus on the results from CLIP-img because encoding models using CLIP-img yield the numerically higher prediction than any of the layers in ResNet-50 (see Figure S2).

**Figure 1:**
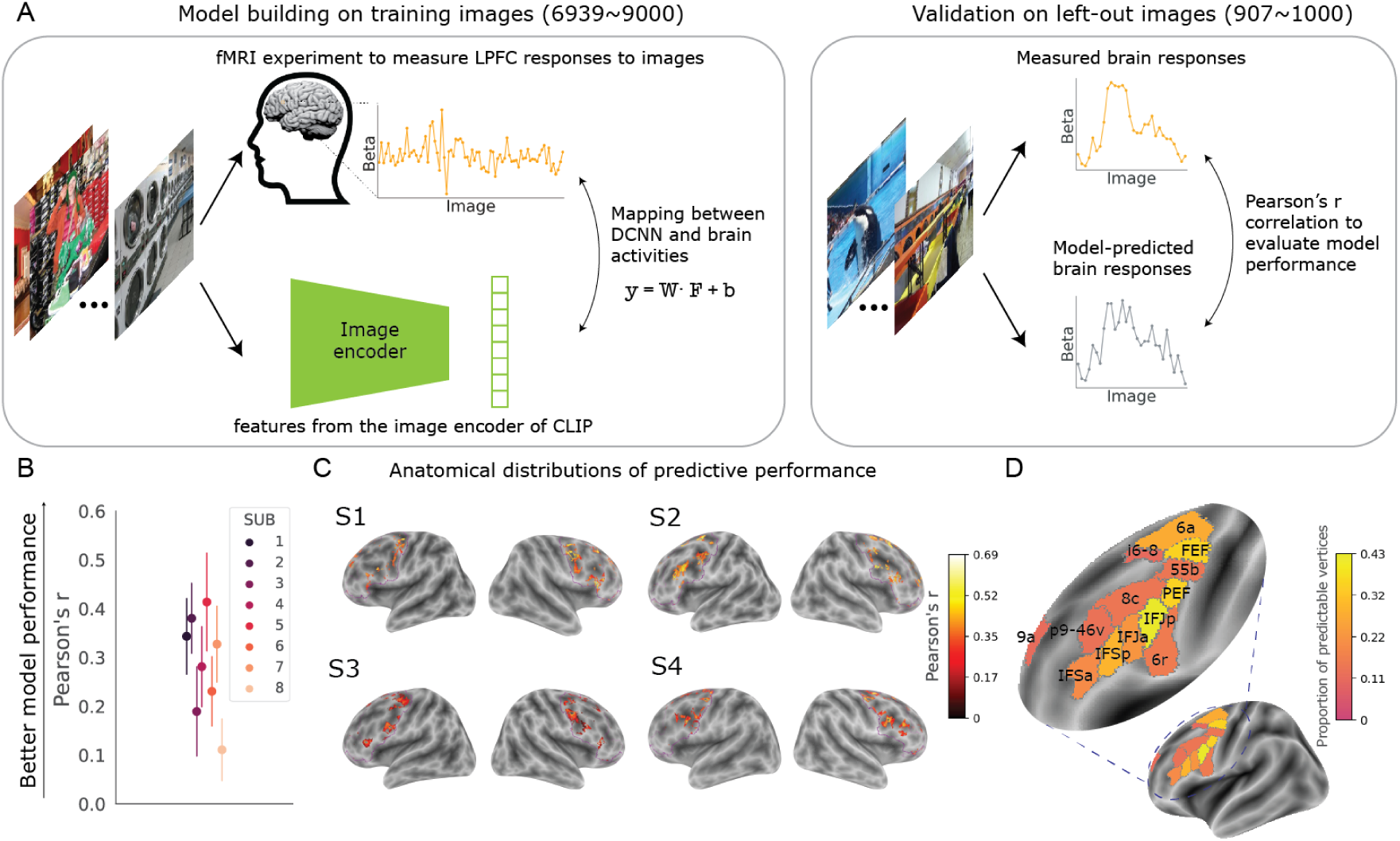
Encoding models robustly predict visual responses in patches of LPFC. (A) Schematic of the cross-validated prediction pipeline. (B) Average Pearson’s r between predicted and observed responses in the top 10% LPFC vertices in the held-out data. Each dot represents a subject. Error bars represent +/- 1 SD. (C) Anatomical distributions of the predictive performance in four example subjects (see Figure S3 for the remaining 4 subjects). The contour of the LPFC mask is marked in purple. (D) The parcels with visually-sensitive vertices (average proportion thresholded at 10%). Parcel labels are defined by [38]

We divided the images into a training set and a test set. Separately for each vertex in a liberal LPFC mask defined based on [38] (see the purple contours in Figure 1C and Figure S1A), we built a ridge regression model to map the CLIP-img features to the brain responses (i.e., single-trial beta estimates) evoked by the training images. We reasoned that only a small proportion of the liberally defined LPFC mask would show sensitivity to visual information and therefore only retained the vertices where the explained variance from the regression models within the training set was in the top 10% among all the LPFC vertices. For each of these selected vertices, we then fixed the regression weights and tested the performance of the encoding model in the held-out images. Model performance was assessed as the Pearson’s r between predicted vs. observed brain responses to the held-out images.

Across all 8 subjects, we observed robust prediction of LPFC in the held-out data (Figure 1B; all ps < .001 based on 10,000 bootstrapping iterations within each subject). Figure 1C shows the anatomical distribution of prediction performance from 4 example subjects (see Figure S3 for the remaining 4 subjects). We also conducted similar analyses using the memory responses to the images to make sure we are not simply identifying vertices related to memory processing (Figure S5).

To further investigate the anatomical location of visually sensitive vertices, within each parcel defined by [38], we calculated the proportion of bilateral vertices that showed cross-validated Pearson’s r > 0.1. This was done separately within each subject. Figure 1D shows the 13 parcels with average proportion of visually-sensitive vertices> 10%. Consistent with findings from non-human primates (e.g., [20, 21, 24]), we also identified visually-sensitive regions in FEF (frontal eye field) and vlPFC (ventrolateral PFC).

### 2.2 Visually-sensitive LPFC vertices show higher functional coupling with visual regions during rest

We next ask: is there also more interaction between these visually-sensitive patches in LPFC and visual regions, compared to the non-visually sensitive parts of LPFC? If these patches genuinely reflect perceptual processing, rather than some artifacts of the computational analysis, we expect this to be the case. To this end, we analyzed the resting-state data included in NSD. Because the LPFC mask we used was intentionally large and liberally defined, we restricted this analysis to vertices belonging to LPFC parcels with over 10% of vertices with a cross-validated Pearson’s r > 0.1), in order to roughly match the anatomical locations of visually-sensitive vs. -insensitive vertices. Specifically, we calculated the functional connectivity between the time series of activity in each vertex of the included LPFC parcels (separately defined for each subject) and the average time series of each of the 68 visual ROIs (see Methods for details on how these visual ROIs are defined).

Figure 2A shows the anatomical distribution of normalized Fisher’s z-transformed correlation values for the LPFC vertices in 4 example subjects (see Figure S4 for the remaining 4 subjects). We then compared the average functional connectivity with visual ROIs for visually-sensitive vs. visually-insensitive (classified based on whether cross-validated Pearson’s r > 0.1) and observed that the visually-sensitive LPFC vertices tend to be more functionally coupled with visual regions during rest compared to those that are spatially close but not predictable based visual features (Fig. 2B; Wilcoxon signed-rank test: T = 0, p = .012).

**Figure 2:**
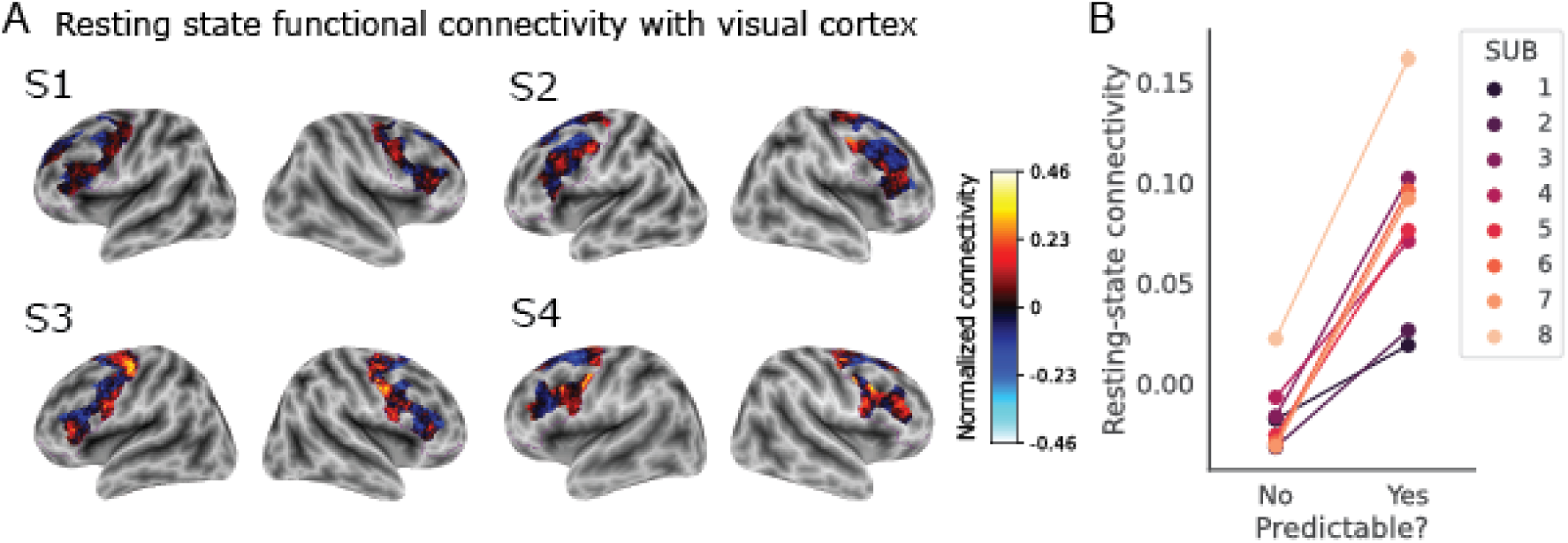
Vertices that are predictable from visual features show higher functional connectivity with visual regions during resting state. (A) Anatomical distributions of the normalized resting state functional connectivity values between LPFC vertices and visual regions in 4 example subjects (see Figure S4 for the remaining 4 subjects). The contour of the LPFC mask is marked in purple. Because the initial LPFC mask is defined liberally as intended, here only parcels (defined by [38]) in which over 10% of the vertices have a cross-validated Pearson’s r > 0.1 are shown, to ensure that the non-predictable vertices are not just entirely outside of visually responsive regions. (B) Average resting-state functional connectivity with the visual cortex in predictable vs. unpredictable vertices in the same parcel defined by [38]. Each line represents a subject. Predictability is defined as having a cross-validated r > 0.1.

### 2.3 The coding scheme of LPFC is more variable across individuals

Although recent work using the same dataset and whole-brain encoding models also presented results showing predictable vertices in frontal regions (e.g., [29]), authors tend to focus on the visual regions. Using neural data of much higher precision, others have studied coding in the LPFC of individual nonhuman primates [21]. Here, we capitalized on the fact that our dataset involves 8 different human subjects, each highly sampled with fMRI (30 to 40 hours of data per subject) to produce robust within-subject measurements. This allowed us to address potential individual differences across subjects.

Specifically, we asked how coding schemes in the visually-sensitive regions in LPFC may differ from that of the ventral visual regions. Given the functional coupling results above, it’s possible that LPFC simply inherits the coding scheme of ventral visual region. Alternatively, LPFC could transform the information passed on to it from the ventral visual region, and such transformation may be specific to each individual.

To compare the coding scheme of PFC to that of ventral visual regions, we focused on the set of images that were seen three times by all 8 subjects (515 unique images in total). Within each ROI of each subject, we calculated the pairwise euclidean distances between the averaged patterns of activities for these images, using only the vertices with cross-validated Pearson’s r > 0.1. We first compared the similarity (measured with Spearman’s rank correlation) between the representational structure (i.e. sets of pairwise activity pattern similarities for all images) in LPFC and that in the ventral visual region within the same subjects to verified that the similarity structure between these images is preserved to some degree within the same individual (see Figure S6), corroborating the results above showing a higher functional coupling between visually sensitive LPFC vertices and visual regions during resting state. We then compared the similarity (also measured with Spearman’s rank correlation) of these representational structures between pairs of subjects within each ROI (see Fig 3A). As shown in Figure 3, we found that the representational structures of these images showed lower across-subject consistency in LPFC than in the ventral visual regions (Wilcoxon’s rank test: T = 0, p < .001).

**Figure 3:**
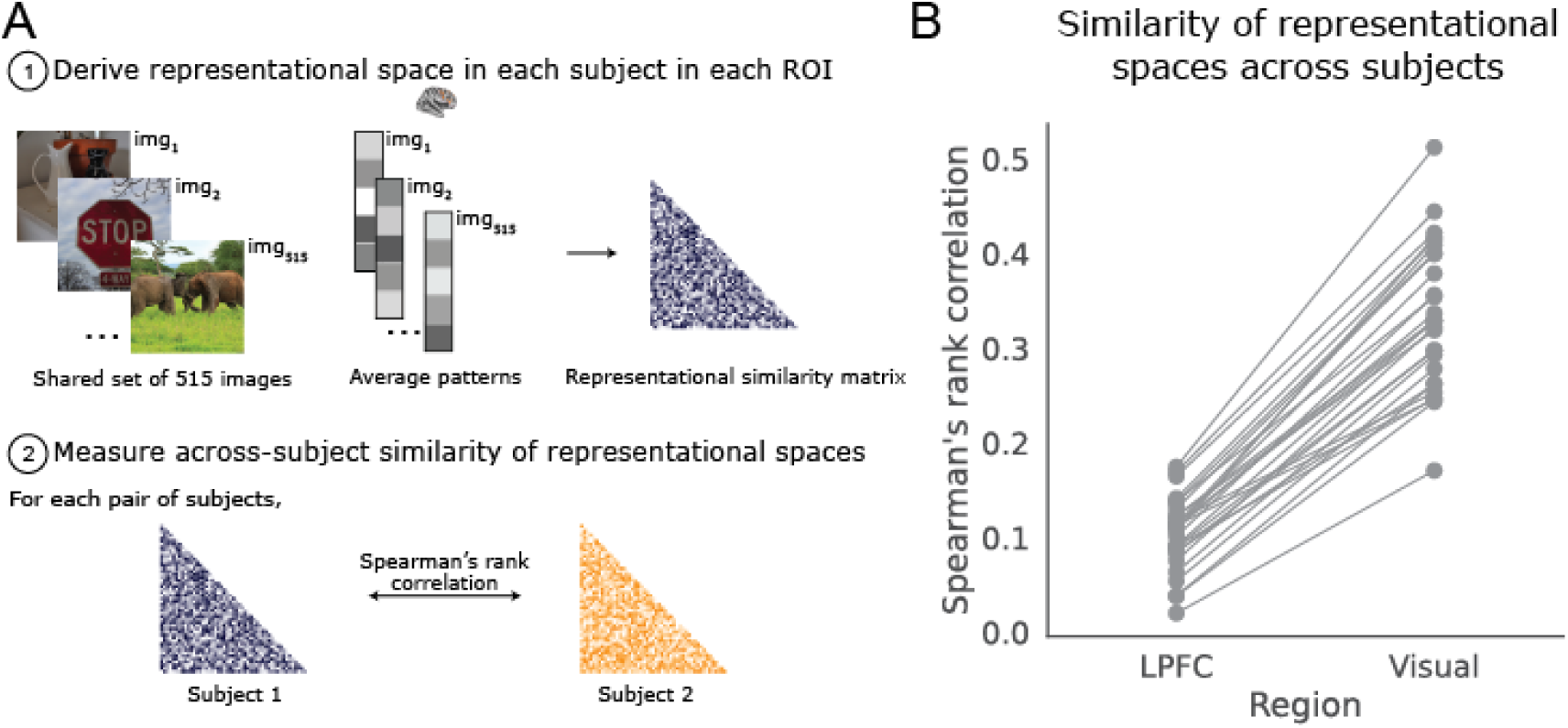
Individual differences in multivariate coding of visual features. (A) Schematic of comparing the representational spaces across subjects. In both LPFC and ventral visual regions, only visually-sensitive vertices (i.e., with cross-validated r > 0.1) are included in the screening. (B) Across-subject similarity of the representational structures in LPFC vs. ventral visual regions. Each line represents a pair of subjects.

To make sure that this is not simply a result of the higher predictability and larger size of the ventral visual regions than LPFC, we subsampled vertices in ventral visual regions to match the number of included vertices as well as the range and mean of cross-validated Pearson’s r values in LPFC (i.e., mean difference in cross-validated Pearson’s r between the subsampled visual vertices and LPFC < 0.06 within each subject). The observed pattern of lower across-subject consistency in LPFC than in the ventral visual regions still holds after the subsampling (p < .001 based on 1000 subsampling iterations; see Figure S7A).

In addition to the individual differences observed in multivariate responses, inspired by recent work using deep-neural-network-based encoding models to identify images that evoke larger responses in category-selective visual regions [39], we screened all 73000 images used as stimuli in the NSD dataset through our robust encoding models (Figure 4A). Shown in Figure 4B are the 8 images predicted to evoke the largest overall averaged responses in the LPFC of the 4 representative subjects (see Figure S8 for the remaining 4 subjects). Corroborating the representational structure results, inspection of these LPFC-activating images reveals a striking variance across individuals: each subject’s LPFC seems to overall ‘prefer’ a different kind of stimuli. To quantitatively confirm the above observations, we correlated the predicted responses for all 73000 NSD images in LPFC across individuals. For comparison, we conducted the same analyses based on the ventral visual regions (see Figure S9 for the 8 images predicted to evoke the largest overall averaged responses in the LPFC of all 8 subjects). Indeed, the correlations of the predicted responses across individuals are lower in LPFC compared to ventral visual regions (Wilcoxon’s rank test: T = 38, p < .001). These results also hold when we subsampled ventral visual vertices to match the average predictive performance to LPFC (p < .001 based on 1000 subsampling iterations; see Figure S7B).

**Figure 4:**
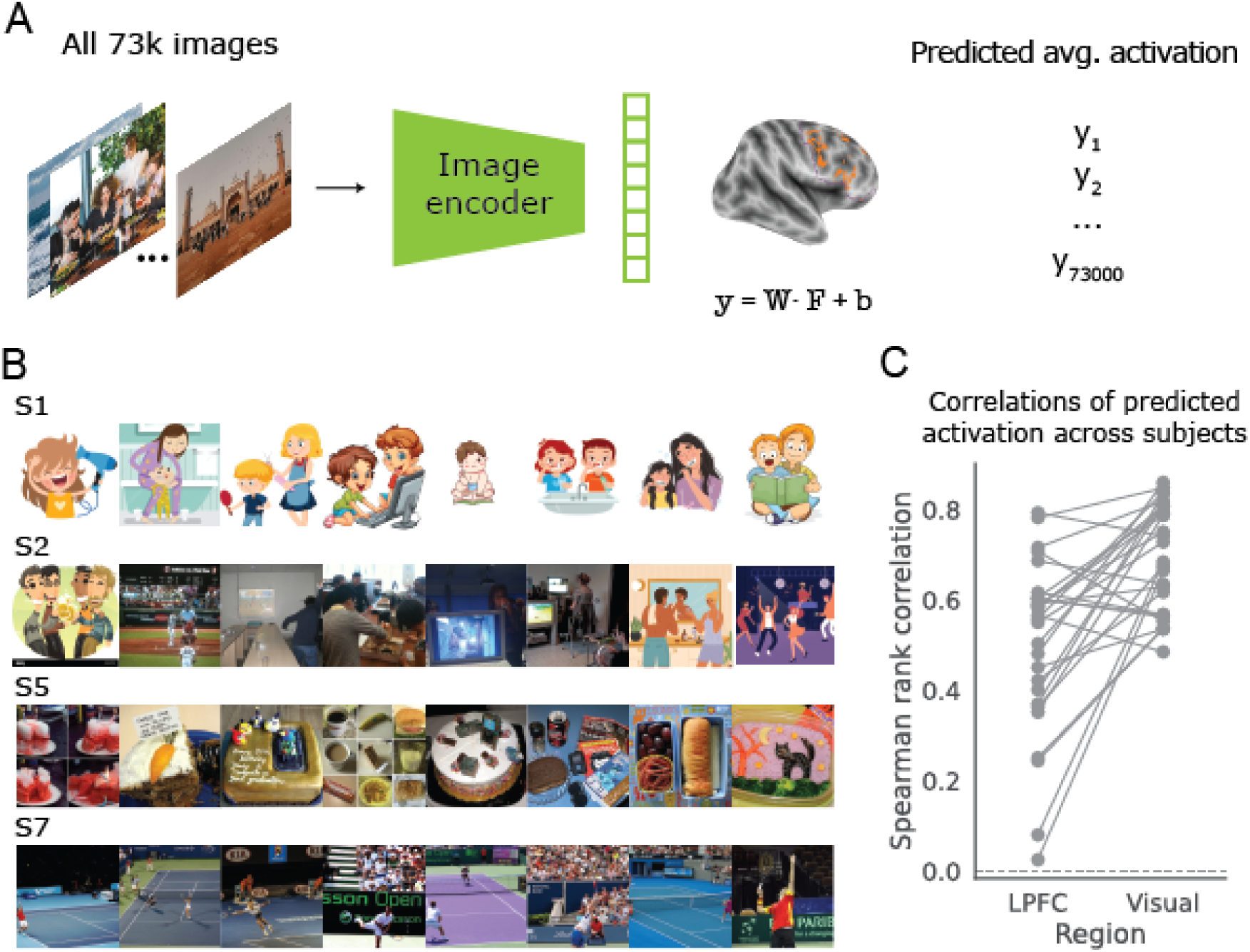
Individual differences in univariate coding of visual features. (A) Schematic of the screening for optimal images to activate the visually-sensitive vertices in a given ROI overall. Here we show an example of the screening for LPFC but the screening is done for both LPFC and ventral visual regions. In both regions, only visually-sensitive vertices (i.e., with cross-validated r > 0.1) are included in the screening. (B) The 8 images with the largest predicted LPFC responses in 4 representative subjects. Images with recognizable human faces have been replaced by cartoons of similar content to comply with the policy of the bioRxiv server. (C) Across-subject correlations of predicted responses in LPFC and ventral visual regions.

### 2.4 Amplified individual differences could result from largely random projection

We show above that the coding scheme of LPFC shows more individual differences compared to that of the ventral visual region, in terms of both multivariate and univariate responses. But what could be the explanation? We attempted to address this question through computational modeling. We turned to a recent model of working memory [40]. The model architecture (see Figure 5A) consists of two layers. The first sensory layer comes with structured coding of a stimulus space (e.g., a color ring) such that neurons are organized topographically according to their selectivity. The second layer is unstructured and has random and reciprocal connections with each of the sensory subnetworks.

**Figure 5:**
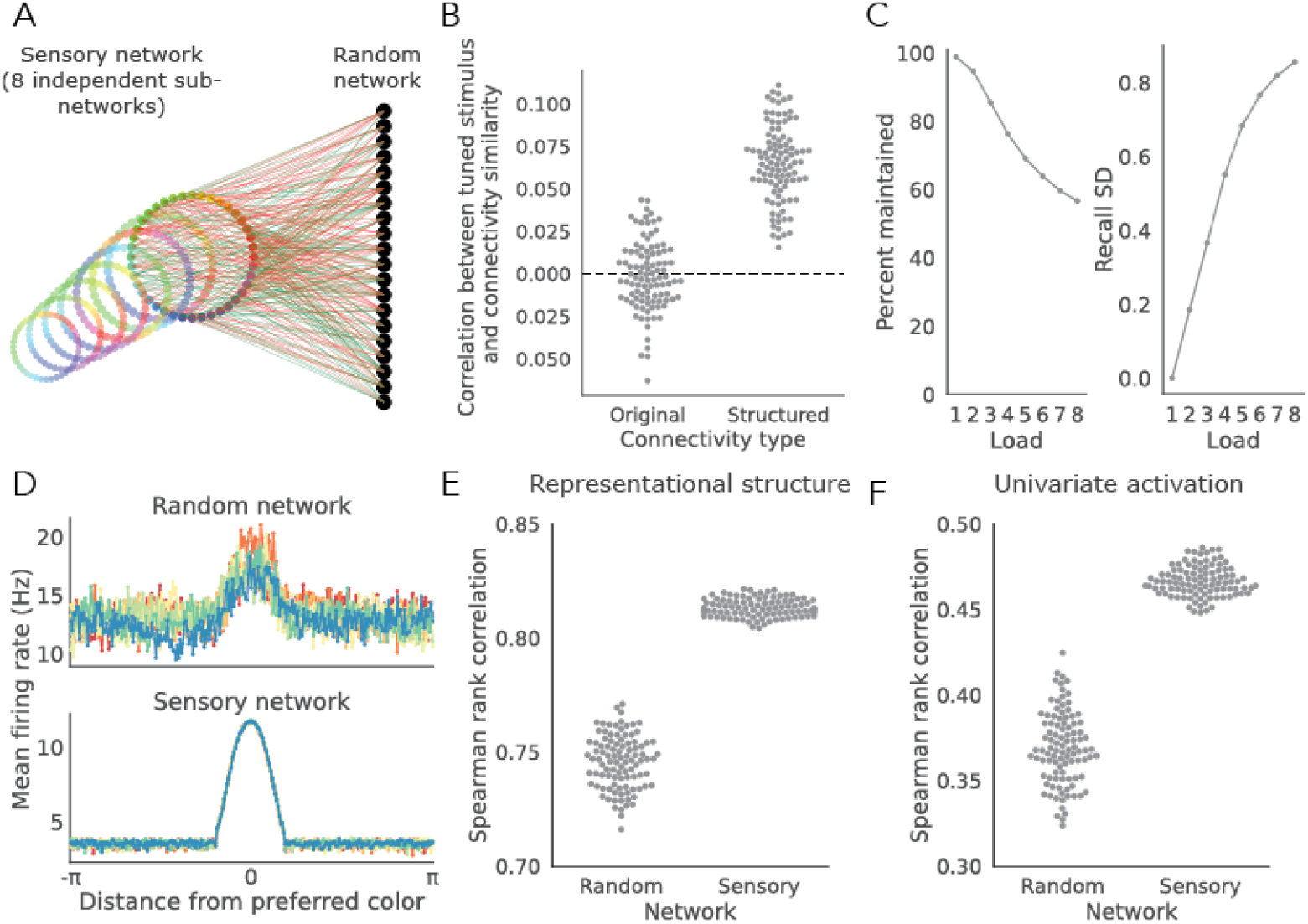
With minimal structure and coding bias, a computational model of working memory [40] exhibits coding characteristics qualitatively similar to our empirical observations. (A) Schematic of the model architecture. The model used in the current study is modified from [40] with the two main adjustments: first, the sensory network demonstrates a bias such that it responds more strongly to a certain stimulus location; second, the connections from sensory network to the random network are not totally random such that 2% of the connections are projecting from sensory neurons tuned to the preferred sensory location to the same set of neurons in the random network. (B) Compared to the original architecture, the average correlation between the similarity of connectivity patterns to the random network and the distance of tuned stimulus locations between pairs of sensory neurons is higher in the modified architecture (Mann Whitney U = 144, p < .001). Each dot represents the correlation between the similarity of connectivity patterns to the random network and the distance of tuned stimulus locations averaged across all pairs of sensory neurons within the same sensory network and all 8 sensory networks from one simulation. (C) The modified model still demonstrates the expected decrease in working memory performance as the number of items held in WM increase, both in terms of the percentage of memory maintained (left) and the precision of the recalled memory (measured as the standard deviation of the recalled error; right). (D) Average firing rates to different stimuli (centered on the preferred stimulus [i.e., the location with larger input strength]) in the random (top) and sensory (bottom) networks. Each color represents a different simulation (with the same preferred stimulus location). There are 8 simulations (matching the number of NSD subjects) in total. (E) Similarity of representational structure across simulated ‘subjects’ in the random vs. sensory network. Each dot is the average Spearman’s rank correlation of the representational structures between pairs of simulated ‘subjects’ for a given preferred stimulus location. (F) Similarity of activation profiles across simulated ‘subjects’ in the sensory network vs the random network. Each dot is the average Spearman’s rank correlation of the activation profile between pairs of simulated ‘subjects’ for a given preferred stimulus location. We simulated 100 preferred stimulus locations (8 different simulated subjects for each location).

There are 8 independent sensory layers/subnetworks, each consisting of 512 neurons connected to the second layer of 1024 neurons through random recurrent projections. As a proof of concept, we decided to explore this model, because the architecture is simple and general. And yet, the network can maintain arbitrary information in memory over time, and recapitulate hallmark behavioral and neural signatures of working memory [40]. Especially relevant to the current study, activities of the random network demonstrate characteristics reminiscent of the prefrontal cortex such as conjunctive coding and elevated activity as memory load increases. Therefore, it allows us to explore the coding relationship between the prefrontal cortex and the sensory cortex.

We modified the original model in two important ways: first, the sensory network demonstrates a bias such that it responds more strongly to a certain stimulus location; this modification was made to mimic the selectivity often observed in the sensory cortex in humans in earlier studies (e.g., FFA [41]) as well as in our study. If there were no differences in the mean population magnitudes between different stimuli, assessing the stability of the univariate responses across individuals is moot. Second, the connections from sensory network to the random network are not totally random such that 2% of the connections are projecting from sensory neurons tuned to the preferred sensory location to the same set of neurons in the random network (see Methods for the detailed implementation). Without this second modification, we could also obtain a qualitatively similar pattern of findings (see Figure S10). However, in this fully random setup, the consistency in univariate activation of the random network across simulated ‘subjects’ is close to zero (see Figure S10C), unlike what we observed empirically in Figure 4C. Therefore, we reason that it is probably unrealistic to assume that the projections from sensory to prefrontal cortex would be truly totally random.

We first verified that the modified architecture is indeed slightly more ‘structured’ (i.e. less random) compared to the original architecture. To this end, for pairs of sensory neurons, we correlated the distance between their selective stimulus locations and the connectivity patterns of their projections to the 1024 neurons in the second, random layer. A high correlation would suggest that neurons coding for similar stimuli also have more similar connectivity patterns to the random layer. As shown in Figure 5B, across 100 simulations of the original and modified architectures respectively, overall the modified architecture showed slightly higher degree of structure compared to the original architecture ((Mann Whitney U = 144, p < .001) while its degree of structure is still very low level (Mean correlation = .06). In addition, we verified that with these modifications, the model still captures classic working memory phenomena such that the percentage of items maintained decreases as the load increases, suggesting limiting capacity, and that the remembered items are less precise (Figure 5C).

After these basic ‘sanity’ checks, we next explored the coding characteristics in the two layers of the network. Figure 5D shows the average firing rates to different stimuli in the sensory layer and random layer respectively for 8 different instances of simulations with the same biased stimulus location. Inspection of this figure suggests that the activation profiles to different stimuli in the sensory layer are very consistent (as indicated by the almost perfectly overlapping curves) whereas the activation profiles in the random network are much more variable. To quantify this observation, we ran 100 simulations, each with a different biased stimulus location in sensory coding. For each biased stimulus location, we ran the simulation 8 different times, matching the number of subjects of NSD, and recorded the responses to 256 different stimulus locations, evenly tiling the entire circular stimulus space (similar to a color wheel). We then compared both the representational structures and univariate activations to these 256 stimulus locations across the 8 simulated subjects, in the sensory and random layers respectively. Mirroring what we observed empirically using NSD data, there are also more ‘individual differences’ once information is passed from the sensory layer to the random layer, both in terms of multivariate pattern responses (Figure 5E) and univariate activation (Figure 5F). Taken together, these modeling results suggest that the amplified individual differences in LPFC coding we observed could be a result of the computational architecture underlying flexible working memory with largely random projection from the sensory network to a high-dimensional random network akin to the prefrontal cortex.

## 3 Discussion

Capitalizing on the recent developments of computer vision models and large-scale sampling of neural responses to natural images within the same individuals, we build encoding models relating visual features to LPFC activities and present evidence for robust prediction within regions of LPFC based on visual features. These visually-sensitive LPFC regions tend to be more functionally coupled with visual regions during rest. However, these visually-sensitive LPFC regions are not simply inheriting information from visual regions verbatim: by comparing the coding profiles across individuals, we demonstrate that the coding profiles of these LPFC regions are more variable across individuals compared to higher-level visual regions, both in terms of multivariate representational spaces as well as overall univariate activation. Computational modeling suggests that such amplification of individual differences can result from the random projection from sensory cortex to prefrontal cortex – an achitecture that is independently known to have many functional benefits and can produce results similar to empirical observations of working memory [40]. Overall, these results suggest that far from being a content-free region reserved for high-level cognition and decision-making (c.f., [42]), LPFC represents rich perceptual information, may play an important role in visual processing that underlies the idiosyncrasies in how different individuals experience the visual world in their own ways.

### 3.1 Seeing beyond the ventral visual stream

Our finding that the human LPFC shows coding of rich visual features adds to the growing literature exploring the functional role of PFC in perception. *How* does the prefrontal cortex support perception beyond what can already be accomplished by the ventral visual stream? Given its extensive reciprocal connectivities with sensory, motor and subcortical regions [43], many have proposed the PFC’s role as a bridge between perception and thoughts/actions [10, 17]. Work in nonhuman primates focusing on the recurrent circuit connecting the ventral visual stream and ventral LPFC (vlPFC) suggests that feedback signals from LPFC are especially important in visually challenging situations, such as recognizing partially occluded objects [23] and recognizing images that are difficult for purely feedforward convolutional neural networks[24]. Another possibility, which isn’t mutually exclusive with the above, is that the prefrontal cortex supports perception by monitoring the quality of perceptual representations and performing metacognitive judgment [44–47]. Having a module for performing such metacognitive judgment in computer vision models (i.e., being aware of the errors oneself made) may help improve the effectiveness and robustness of such models by reducing the likelihood of hallucination [48]. Our finding of the amplified individual differences in PFC coding may provide a new angle to understanding the functional role of PFC in perception: PFC may turn the stable representational landscape of visual experiences – provided by the ventral visual stream – into a new space that is more tailored to the individual’s goals and priority. These converging lines of evidence suggest that PFC compliments the visual system by improving the robustness, flexibility and adaptiveness of perception and thus leading to representations that are more useful for thinking and acting.

### 3.2 Neural basis for individual differences in perception

The majority of neuroscience research on perception has focused on deriving generic principles on how the brain supports perception by aggregating data from multiple participants and treating individual variability as ‘noise’. In contrast to this tradition, recent work has started to focus on the neural basis of individual differences of perception. Early visual cortex such as V1 has been linked to individual differences in contrast sensitivity [49] and object size perception [50]. Multivariate representational spaces of objects in inferior temporal cortex capture unique aspects of the perceived similarity between objects [51]. Particularly relevant to our study, the representational similarity of morphs between two facial identifies in LPFC, but not in face selective areas in ventral visual stream, is correlated with individual differences in reaction times for those morphs in a visual search task [52]. This seems to corroborate our finding of amplified individual differences in coding schemes of PFC compared to those of the ventral visual stream.

With the relatively large scale of the dataset used in our current study[32], both in terms of stimuli and data quantity, and the use of neural network features in our encoding models, we are able to chart the complexity and richness of the PFC coding schemes across different participants. However, we do not yet know how the neural coding schemes in LPFC contribute to perceptual behavior. One intriguing hypothesis is that the observed individual differences in PFC coding schemes reflect individual differences in internal models of the world (i.e., idiosyncrasies in our expectation of what the world should be like by default) that scaffold our perception (e.g., [53]). In follow-up studies, we are now exploring the behavioral consequences of such individual differences in PFC coding with large-scale psychophysical experiments, using stimuli tailored to each individual participant. We hypothesize that the observed individual differences in PFC coding may reflect *individualized* priority maps [54, 55] such that the LPFC-activating and deactivating items identified separately for each individual should be prioritized accordingly based on this individual’s priority map and therefore we should observe selective behavioral advantages/disadvantages for these items. Moreover, given that we were able to obtain our current results, where individuals were tested over the course of many months, we expect that these priority maps may be relatively stable and possibly not task-dependent.

### 3.3 Sources of individual differences in prefrontal coding

Where do the observed individual differences in the coding schemes of PFC come from? The computational modeling we conducted based on a flexible model of working memory [40] suggests a somewhat provocative interpretation: the observed individual differences in PFC coding could just be a by-product of a neural architecture that is able to flexibly maintain information in working memory with limited capacity. The model fails to generalize to better maintaining stimuli in working memory that it hasn’t encountered before, if the projection from the sensory layer to random layer is trained and optimized [40]. That is, the random projection from the sensory layer to random layer seems critical for the flexibility of the model. Therefore, the observed amplification of individual differences from the sensory cortex to PFC could just be an inevitable result of the random projection where the goal is to maintain information in working memory, rather than to amplify individual differences *per se*.

However, we suspect that this is unlikely to be the full answer. Research strongly highlights the genetic underpinnings of both structural and functional connectivity of PFC (e.g., [56, 57]). It’s possible that genetic factors contribute to the empirically observed individual differences in PFC coding described in our study.

Another intriguing possibility is that the individual differences result from differences in visual ‘diet’ (e.g., [58, 59]). The accumulated visual experiences of distinct individuals shape their internal models of the external world and manifest in the form of a more individual specific representational scheme in the PFC, in contrast to one that’s more or less shared across individuals in the visual regions. These intriguing possibilities can be further explored in twin studies or cross-cultural populations.

### 3.4 Is it really ‘visual’?

One important theoretical question regarding the interpretation of our results is: are the LPFC regions identified by our encoding models truly ‘visual’? Or are they encoding some more abstract and conceptual features that need to be derived from sensory experiences but do not necessarily come from vision? For example, a PFC ensemble encoding the properties corresponding to ‘waterfall’ may respond similarly to an image of waterfall or to a sound clip of waterfall running down. This is a possibility since some neurons in PFC are indeed polysensory such that they respond to both images and sounds (e.g., [21, 60, 61]).

Leveraging the fact that all the NSD stimuli come with human-generated captions from 5 different observers and CLIP model is jointly trained on images-captions pairs and thus also comes with a text encoder, we further explore the representational format in LPFC in a supplemental analysis (see Figure S11). Inspired by recent work using NSD demonstrating the alignment between Large Language Models (LLMs) text embeddings of image captions and activities to natural scenes in high-level visual regions[62], we asked whether LPFC responses to scene images can also be predicted based on embeddings extracted from the CLIP-text encoder for the images captions. Interestingly, such textural embeddings from CLIP also capture substantial variance in LPFC, consistent with recent work from human intracranial recordings demonstrating the alignment between LLM embeddings for description of images and visually-evoked activities in frontoparietal regions[63]. However, unlike in [63] where they found that the representational structures of 28 images of famous places and humans in LPFC are better explained by CLIP-text embeddings than CLIP-image embeddings, we observe a different trend in the NSD dataset where each subject saw around 10000 unique scene images from diverse categories. Specifically, the encoding models based on CLIP-img show a slight advantage across all 8 NSD subjects in terms of predictive performance(see Figure S11B). Along with the pattern observed in Figure S2 where the later layers of ResNet-50 tend to predict LPFC activities better, these results suggest that the representational format in LPFC is indeed richer and more complex, but perhaps not fully distillable only into words.

Given that the stimuli used in the NSD dataset are all static images without sound, we are unable to address whether we would have been able to identify the same set of LPFC regions if we use stimuli from a different modality yet conveying similar abstract information. After all, the NSD subjects still processed the images but not the captions. Future studies can investigate this question by using multi-modal stimuli with rich visual and auditory information.

### 3.5 Methodological implications

Our results also provide several methodological implications for studying the role of LPFC in perception. First, our results on individual differences suggest that to identify visually sensitive regions in LPFC, the ideal analysis may need to be conducted within each individual, given the anatomical variations of those visually-sensitive regions (see Figure 1C) and the substantial individual differences in what visual stimuli tend to drive each person’s LPFC. The use of a common set of stimuli in some previous group studies might have contributed to the impression that visual responses in LPFC are weak or absent.

Second, the finding of amplified individual differences in the representational structures of PFC also has implications for applying methods for aligning different individuals’ brain into the same space such as hyperalignment[64] and shared response modeling[65] to data from LPFC. A fundamental assumption of such methods is that there is a common information space for coding sensory inputs that is shared and consistent across individuals. However, our results suggest that this assumption may be less straightforwardly applicable to the LPFC. Future work should assess whether this complication may necessitate new methods.

In conclusion, our study highlights the underappreciated role of PFC in perception with a specific focus on LPFC and individual differences. The LPFC not only codes rich and complex visual features but may underlie the diversity of how individuals perceptually experience the world. More broadly, our study showcases the power of combining powerful tools from modern machine learning, high-field fMRI and dense sampling (see also [66–68]) to reveal novel aspects of how the human brain perceives and constructs internal representations of the external world.

## 4 Methods

### 4.1 The Natural Scenes Dataset

All the empirical results presented in the current study are based on data from the Natural Scenes Dataset (NSD) [32]. Complete details of the study can be found in the Methods section of [32]. Briefly, there are 8 subjects who viewed 9209 to 10000 unique scene images (each presented up to 3 times) while being scanned at 7T and performing a recognition memory task.

#### Task data preparation

We analyzed the single-trial betas (version 3 [betas fithrf GLMdenoise RR]) in native surface space (resampled from 1-mm subject native volume) derived from all available NSD task sessions, provided as part of the NSD data release. The single-trial betas were normalized across time within each NSD session, separately for each vertex, to account any potential systematic difference in the overall responsiveness in BOLD signals between NSD sessions. An averaged beta response to each image was then derived by averaging over all the normalized betas corresponding to the given image. We included any images shown in our analysis (regardless of the number of repetitions). For Subjects(S) 1, 2, 5 and 7, neural responses to 10000 unique images repeated 3 times over 40 sessions were included. For the remaining subjects, breakdown is as follows (number of unique image [number of sessions]): S3-9411 [32], S4-9209[30], S6-9411[32], S8-9209[30].

#### Resting state data preparation

Although two kinds of resting state runs were collected by NSD (one where subjects were instructed to simply stay awake and fixate and the other where they were additionally instructed to inhale deeply when fixation cross changed to red color), we only analyzed the resting state runs without the breath-holding (func1pt8mm in native volumetric space with 1.8 mm spaital resolution and 1.33 second temporal resolution). To minimize the effect of motion on functional connectivity estimates, we excluded any runs that meet any of the following criteria: 1) mean frame-to-frame displacement > 0.15 mm; 2) maximum frame-to-frame displacement > 2 mm; 3) maximum translation > 2 mm; 4) maximum rotation > 3 degrees. The numbers of runs included after applying these criteria are as follows: S1-18, S2-8, S3-5, S4-9, S5-10, S6-10, S7-10, S8-8. In other words, the amount of resting state data ranges from 25 minutes to 90 minutes.

Additional preprocessing was done using the nilearn.signal.clean function on the resting state data including: exclude the first 4 TRs, detrending, bandpass filtering (range: [0.009, 0.08] Hz), regressing out mean signals from white matter, gray matter, and whole-brain and 6 motion parameters as well as the derivatives for all aforementioned measures, and z-scoring over time. The preprocessed data in volumetric space was then resampled via cubic interpolation onto the subject-native cortical surfaces (which exist at 3 different depths), and averaged across depths to get the resting state timeseries in native surface space. This mapping was done using the python toolbox provided by the NSD team (https://github.com/cvnlab/nsdcode.git).

#### ROI definition

##### Lateral Prefrontal Cortex (LPFC)

We defined an intentionally large LPFC mask based on [38]. The LPFC mask includes parcels with the following labels provided by [38]: FEF, PEF, 55b, 8Av, 8Ad, 9m, 8BL, 9p, 10d, 8C, 44, 45, 47l, a47r, 6r, IFJa, IFJp, IFSp, IFSa, p9-46v, 46, a9-46v, 9-46d, 9a, 10v, a10p, 10pp, 6a, i6-8, s6-8, p10p, p47r. As shown in Figure S1A, the lPFC mask is quite large.

##### Ventral visual stream

For comparison with LPFC coding of visual features, we additionally defined a ventral visual stream mask using the streams ROIs provided by NSD. The ventral ROI was drawn to follow the anterior lingual sulcus (ALS), including the anterior lingual gyrus (ALG) on its inferior border and to follow the inferior lip of the inferior temporal sulcus (ITS) on its superior border. The anterior border was drawn based on the midpoint of the occipital temporal sulcus (OTS). See Figure S1B for the visual ventral stream mask in one example subject.

##### Visual ROIs

To investigate the funcitonal coupling between LPFC and visual regions during resting state, we additionally defined a broad set of visual ROIs separately for each hemisphere in three different ways: 1) for ROIs that can be mapped with a population receptive field (prf) task (such as V1-ventral and V2-ventral), we used the manually-drawn ROIs provided by NSD (the prf-visualrois); 2) for ROIs that can be identified in a functional localizer task (such as the occipital face area), we again used the manually-drawn ROIs provided by NSD (the fLoc-faces/words/places/bodies); 3) the remaining ROIs (such as the intraparietal sulcus) were defined based on a probablisitic atlas ([69]; the Kastner2015 ROIs in NSD). To ensure that we are only including ROIs with meaningful visual signals, we additionally excluded any ROIs where there were fewer than 100 visually responsive vertices in any given subject. Visual responsiveness is defined as > 1% variance explained by a simple ON-OFF GLM, averaged across all NSD sessions. See Table S1 for the full list of the 68 visual ROIs included and the median number of visually-responsive vertices.

### 4.2 Analysis

#### Feature-based encoding models

To build encoding models of LPFC within each subject (see Figure 1A), for the presented images, we first extracted activations from the image encoder of a CLIP (Contrastive Language-Image Pre-Training) network (ViT-B/32 backbone; [36]). We divided the images into a training set (consisting of images seen only by the given subject) and a test set (consisting of images shared across subjects). For S1, 2, 5 and 7, the training set includes 9000 images and the test set includes 1000 images. For the remaining subjects, breakdown is as follows ([number in the training set, number in the test set]): S3[8481, 930], S4[8302, 907], S6[8481, 930], S8[8302, 907].

Separately for each vertex in the LPFC mask (see the purple contours in Figure 1C or Figure S1A), we built a ridge regression model to map the CLIP-img features to the averaged beta estimates evoked by the training images. We reasoned that only a small proportion of vertices inour liberally defined LPFC mask would be visually-sensitive. Therefore, we only retained the top 10% vertices in terms of the explained variance from the regression models within the training set to reduce the amount of computational resources needed. The regularization parameter for each vertex was chosen via 3-fold cross-validation across the training set from a range of 7 different parameters evenly spaced on a log scale from 10*^—^*1 to 10^6^. For each of these selected vertices, we then tested the performance of the ridge regression model in the held-out images. Performance was assessed as the Pearson’s r between predicted vs. observed responses to the held-out images. To assess statistical significance, we ran 10000 bootstrapping iterations by resampling the images in the testing set with replacement.

For comparison, we built encoding models based on vertices in the ventral visual region and evaluated the performance using the same cross-validation pipeline as described above. Because we reasoned that most of the ventral visual region vertices should be visually-sensive, we conducted cross-validation on all the vertices in the ventral visual mask, rather than only the top 10%.

#### Ruling out the effect of memory performance

As LPFC has been found to be involved in successful memory encoding and retrieval ((e.g., [70]), [71], [72]), we wanted to rule out the possibility that the CLIP-feature-based models are simply picking up activatiies related to memory processing per se. To this end, we built models using memory performance instead of CLIP-features as the predictors in the regression models. Results for this analysis are presented in Figure S2.

##### Memory-encoding-based models

To identify LPFC vertices that track successful memory encoding in each subject, we focused on single-trial betas corresponding to the first presentation of images that repeated at least twice and the behavioral performance when the image was presented for the second time. We only considered whether the image was presented for the second time but not the third time because this would be more akin to how subsequence memory effect is usually measured in the literature (reviewed in [73]). In addition to whether the subject correctly responded ‘old’ when the image was repeated for the second time, we also included the delay between the first and second presentations of the images as an additional regressor given the variability in the delays between the first and sceond presentations in NSD (see Extended Data Figure 1a in [32]). As the CLIP-feature-based models, we trained the models on the brain activities and memory performance corresponding to the images seen only by the given subject and evaluated them on those corresponding to images shared across subjects. Performance was assessed as the Pearson’s r between predicted vs. observed brain responses to the held-out images.

##### Memory-retrieval-based models

To identify LPFC vertices that track successful memory retrieval in each subject, we focused on single-trial betas corresponding to the second and/or third presentation of images that repeated at least twice and the behavioral performance when the image was repeated. In addition to whether the subject correctly responded ‘old’ when the image was repeated, we also included the delay between the current and last presentations of the images as an additional regressor. As the CLIP-feature-based models, we trained the models on the brain activities and memory performance corresponding to the images seen only by the given subject and evaluated them on those corresponding to images shared across subjects. Performance was assessed as the Pearson’s r between predicted vs. observed brain responses to the held-out images.

#### Resting state functional connectivity

The resting state functional connectivity analysis was limited to LPFC parcels (defined separately within each subject) with more than 10% vertices showing cross-validated Pearson’s r > 0.1. Separately for each subject and resting state run, we calculated the correlation (Fisher z-transformed Pearson’s r) between the timeseries of each vertex in the included LPFC parcels and the average timeseries of each of the 68 visual ROIs shown in Table S1. The resulting correlation values were further z-scored within each visual ROI to equate for baseline differences in overall connectivity with LPFC across the various visual ROIs. Finally, we averaged the z-scored correlation values across all visual ROIs and all available resting state runs to obtain an average resting state connectivity measure with visual ROIs for each LPFC vertex. Finally, we compared the average connectivity values of visually-sensitive LFPC vertices to those that are visually-insensitive across the 8 subjects with Wilcoxon signed-rank test.

#### Comparing the coding schemes in LPFC vs. ventral visual regions

To compare the coding schemes in LPFC and ventral visual regions, we adopted both multivariate and univariate approaches, limiting our analyses to only vertices with a cross-validated Pearson’s r > 0.1 in both regions.

##### Representational similarity

To compare the representational structures in LPFC/visual regions across subjects, we focused on the set of 512 images seen three times by all subjects. For each ROI of each subject, we derived a response pattern for each image by averaging over the three repetitions. We calculated the pairwise euclidean distances between the image-specific response patterns to obtain the representational structure of the ROI in the given subject. We then compared the similarity (assess via Spearman’s rank correlation) of these representational structures in each ROI for each pair of subjects. To assess the statistical significance of the difference in the two ROIs’ across-subject similarities of representational structures, we used Wilcoxon’s signed-rank test.

##### Predicted univariate activation

We built subject-specific encoding models to predict the average responses in the predictable vertices of each ROI. We then screened all 73000 NSD images through these encoding models of LPFC/visual region activity. To quantify individual differences, we calculated the Spearman’s rank correlation of the predicted responses for all images for pairs of subjects with Spearman’s rank correlation.

##### Subsampling analysis

To rule out effects from the ROI size and overall predictability, we repeated the above two analyses of representantional similarity and univariate activation on subsampled vertices in the ventral visual regions. Specifically, within each subject we matched the number of predictable vertices and the range and overall predictability to LPFC(i.e., mean difference in cross-validated Pearson’s r between subsampled ventral visual vertices and LPFC vertices < 0.1). We repeated the subsampling 1000 times to assess the stability of the results.

### 4.3 Computational modeling

To model the interaction between sensory cortex and prefrontal cortex, we made slight modifications to a previously established model of flexible working memory [40]. For details of the model setup and parameters, please refer to the Method Details section of [40].

Briefly, the model is a two-layer network of Poisson spiking neurons. The first layer is a sensory network consisting of 8 ring-like subnetworks. Each subnetwork comprises of N*_Sensory_* = 512 sensory neurons. Every neuron *i* in a subnetwork has a associated tuning angle of Φ*_i_* = 2π*i/N_Sensory_*. Each sensory subnetwork can be used to encode a continuous feature space (e.g., a color ring as shown in Figure 5A). Each sub-network receives sensory input independently. The second layer is a random network comprising of *N_Random_* = 1024 neurons randomly connected to each oof the 8 simulated sensory sub-networks. We have kept the parameters exactly the same as in [40] except in the following two aspects.

#### Response selectivity in the sensory network

To mimic the selectivity often observed in the sensory cortex in humans in earlier studies (e.g., FFA [41]) as well as in our study, we made the sensory network respond stronger to a certain stimulus location. We achieved this goal by increasing the strength of external sensory input at the biased stimulus relative to the non-biased stimuli whereas the original model has the same strength input strength for all stimuli. Specifically, the sensory input to neuron i is

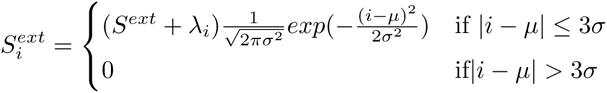

where µ is the center of the Gaussian inputs, and σ is the width of the input. The same as in the original model, σ was defined as a fraction of the total number of neurons in the sensory sub-network, σ = *N_sensory_*/32, which was 16 neurons for the presented network. The same as in [40], S*^ext^* is set to 10. λ*_i_* is an additive modulation factor that we introduce to make the strength of external sensory input depend on the tuned stimulus of neuron i. Specifically, λ*_i_* is

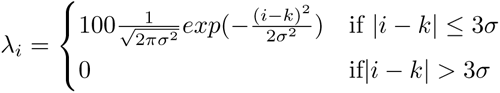

where *k* is the index of the preferred stimulus that ranges from [1, N*_sensory_*] and stays consistent for each simulation.

Minimal structure in the projection from sensory to random layers

To inject a little structure into the projection from sensory to random layers, we re-arrange a subset of the connections between sensory and random layers. For each neuron in the random layers, there is a 10% probability that the connections to and from this neuron would be rearranged to inherit some structures. Specifically, for each of these random neurons with re-arrangement, we swap the connections between this random neuron and each sensory network such that 7 sensory neurons whose indices falling within [*k — σ, k + σ*] (with our current set of parameters: [*k* — 16, *k* + 16]; thus about 20% of the neurons within this range) project to this random neuron. We achieve this by sampling 7 neurons without replacement from the indices range [*k — σ, k + σ*], where the probability of being selected is a Gaussian centering around k such that neurons for which the tuning is closer to k (i.e., the preferred stimulus) have a higher likelihood of being selected to project to this random neuron. To maintain the number of connections consistent with the fully random architecture, in each of this swapping iteration, we randomly eliminate 7 other pairs of connections between this random neuron and neurons tuned to stimuli outside of [*k — σ, k + σ*]. Effectively, this procedure should result in 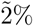 of the connections between sensory and random layers going from sensory neurons falling within the preferred range [*k — σ, k + σ*] to the same set of random neurons.

To verify that this procedure is indeed successful in injecting structure in the connections, we simulated 100 networks using the original (thus fully random) setup by [40] and another 100 networks using the modified setup described above. For each network, we first calculate the absolute distances between the tuned stimuli for pairs of sensory neurons within each sensory subnetwork. We then calculate the euclidean distances of connectivity patterns from pairs of sensory neurons within each sensory subnetwork to the random layer. For all pairs of sensory neurons in the same sensory subnetwork, we correlate the absolute distances in tuning and the euclidean distances of connectivity patterns. Lastly, we average the correlations across 8 sensory subnetworks to obtain an overall measure how related the connectivity patterns from sensory neurons to random neurons are to the similarity of tuning, as a measure of the degree of structure in the sensory-random connectivity for each simulated network.

#### Simulation for exploring individual differences

To explore individual differences in the coding of sensory and random networks, we ran 100 simulation groups, each with a different biased stimulus location in sensory coding. For each biased stimulus location, we ran the simulation 8 different times, matching the number of subjects of NSD.

##### Model working memory performance

To make sure that the modified architecture still allows for the maintenance of information in working memory (WM) and demonstrates the canonical WM phenomena, we presented 768 WM trials to each simulated network and recorded the model’s performance. The WM load ranges from 1 to 8, chosen randomly on each trial. The input center (i.e., the color to be held in mind) is also randomly chosen for each subnetwork receiving input. Each stimulus is presented for 100 ms followed by a delay period of 1 second. Similar to [40], we calculated the percentage of memory maintained as proportion of successfully maintained memories among initial inputs to sensory sub-networks. Successful maintenance is defined as the norm of the activity vector for a given sub-network exceeding 3Hz. In addition, we calculate the precision of WM recall as the standard deviation of drifts of recalled memory. The drift of a memory is simply calculated as the absolute value of the angle distance between the initial input value and the decoded value at the end of the memory delay. These measures were calculated on every trial and then summarized by binning them into different loads.

##### Coding in sensory and random layers

To explore the coding of different stimuli, we additionally presented another 768 trials to each network. The WM load is always 1 and the subnetwork receiving input is always subnetwork 1 on each trial. To make sure we measure responses to the entire stimulus space, we present 256 different stimuli, evenly spaced on the stimulus ring and covering the entire ring. Each stimulus is presented 3 times (mimicking the NSD design). To record the activity evoked by the different stimuli, we take the activities in the first subnetwork of the sensory layer at 100 ms after stimulus onset and the activities in the random layer at 200 ms after the stimulus onset.

To explore consistency of univariate activation across the simulated ‘subjects’, we calculate the pairwise Spearman’s rank correlation of the average population responses to the 256 stimuli between pairs of simulated ‘subjects’, separately for the sensory (only subnetwork 1) and random layers. This is done within all possible pairs of 8 simulations with the same preferred stimulus k. We repeat this process for the 100 groups of 8 simulations difference preferred stimulus.

To explore the consistency of multivariate patterns, we first derive the representational similarity matrices for the 256 stimuli by calculating the euclidean distances between the response patterns corresponding to pairs of stimuli for each simulated ‘subject’, separately for the sensory (only sub-network 1) and random layers. We then calculate the pairwise Spearman’s rank correlation of the representational similarity matrices between pairs of simulated ‘subjects’, separately for the two layers. This is done within all possible pairs of 8 simulations with the same preferred stimulus k. We repeat this process for the 100 groups of 8 simulations difference preferred stimulus.

## Acknowledgments

We are grateful for the RIKEN Information Systems Division for providing and maintaining the high performance computing cluster (Hokusai) utilized by this study. We would like to thank Tim Buschman and Sebastian Eydam for helpful discussion on modifying the computational model of working memory.

## Code and Data Availability

All data is made publicly available via the Natural Scenes Dataset (https://naturalscenesdataset.org/). Analysis code is available on Github (https://github.com/Qi-Lin7/NSD PFC.git).

---

**Table S1:**
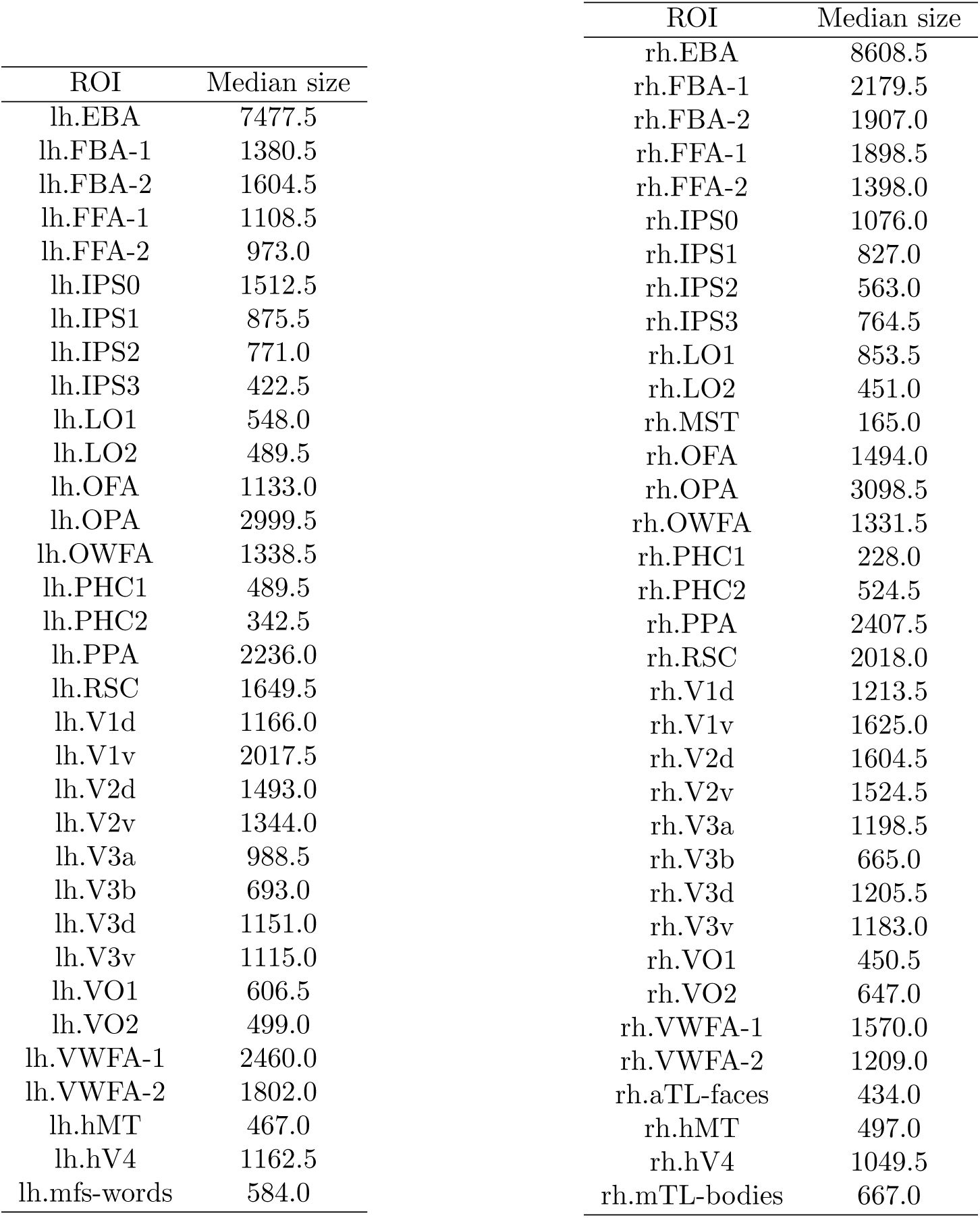
Visual ROIs in the resting state analysis.

| ROI | Median size |
| --- | --- |
| lh.EBA | 7477.5 |
| lh.FBA-1 | 1380.5 |
| lh.FBA-2 | 1604.5 |
| lh.FFA-1 | 1108.5 |
| lh.FFA-2 | 973.0 |
| lh.IPS0 | 1512.5 |
| lh.IPS1 | 875.5 |
| lh.IPS2 | 771.0 |
| lh.IPS3 | 422.5 |
| lh.LO1 | 548.0 |
| lh.LO2 | 489.5 |
| lh.OFA | 1133.0 |
| lh.OPA | 2999.5 |
| lh.OWFA | 1338.5 |
| lh.PHC1 | 489.5 |
| lh.PHC2 | 342.5 |
| lh.PPA | 2236.0 |
| lh.RSC | 1649.5 |
| lh.V1d | 1166.0 |
| lh.V1v | 2017.5 |
| lh.V2d | 1493.0 |
| lh.V2v | 1344.0 |
| lh.V3a | 988.5 |
| lh.V3b | 693.0 |
| lh.V3d | 1151.0 |
| lh.V3v | 1115.0 |
| lh.VO1 | 606.5 |
| lh.VO2 | 499.0 |
| lh.VWFA-1 | 2460.0 |
| lh.VWFA-2 | 1802.0 |
| lh.hMT | 467.0 |
| lh.hV4 | 1162.5 |
| lh.mfs-words | 584.0 |
| rh.EBA | 8608.5 |
| rh.FBA-1 | 2179.5 |
| rh.FBA-2 | 1907.0 |
| rh.FFA-1 | 1898.5 |
| rh.FFA-2 | 1398.0 |
| rh.IPS0 | 1076.0 |
| rh.IPS1 | 827.0 |
| rh.IPS2 | 563.0 |
| rh.IPS3 | 764.5 |
| rh.LO1 | 853.5 |
| rh.LO2 | 451.0 |
| rh.MST | 165.0 |
| rh.OFA | 1494.0 |
| rh.OPA | 3098.5 |
| rh.OWFA | 1331.5 |
| rh.PHC1 | 228.0 |
| rh.PHC2 | 524.5 |
| rh.PPA | 2407.5 |
| rh.RSC | 2018.0 |
| rh.V1d | 1213.5 |
| rh.V1v | 1625.0 |
| rh.V2d | 1604.5 |
| rh.V2v | 1524.5 |
| rh.V3a | 1198.5 |
| rh.V3b | 665.0 |
| rh.V3d | 1205.5 |
| rh.V3v | 1183.0 |
| rh.VO1 | 450.5 |
| rh.VO2 | 647.0 |
| rh.VWFA-1 | 1570.0 |
| rh.VWFA-2 | 1209.0 |
| rh.aTL-faces | 434.0 |
| rh.hMT | 497.0 |
| rh.hV4 | 1049.5 |
| rh.mTL-bodies | 667.0 |

**Figure S1:**
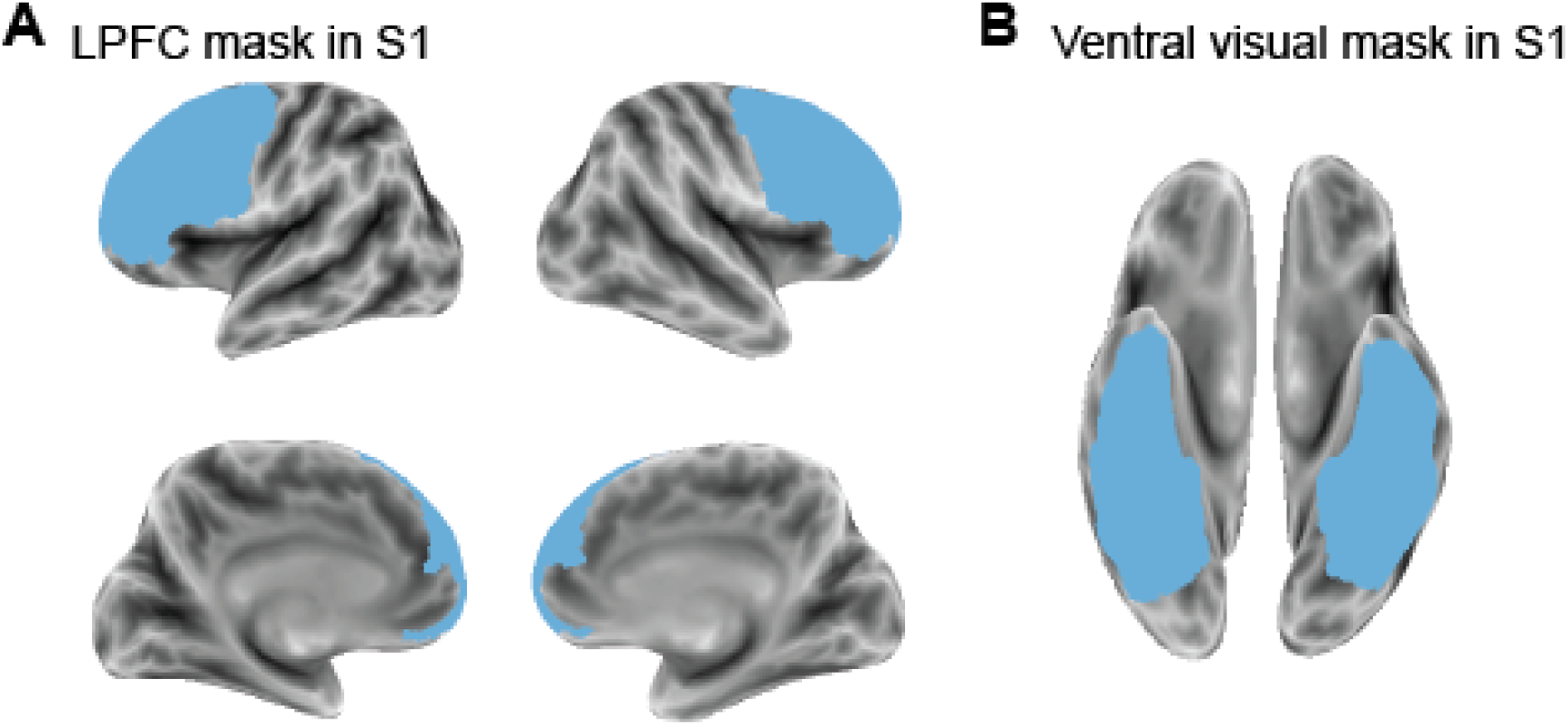
The LPFC(lateral prefrontal cortex) and ventral visual stream masks in one example subject.

**Figure S2:**
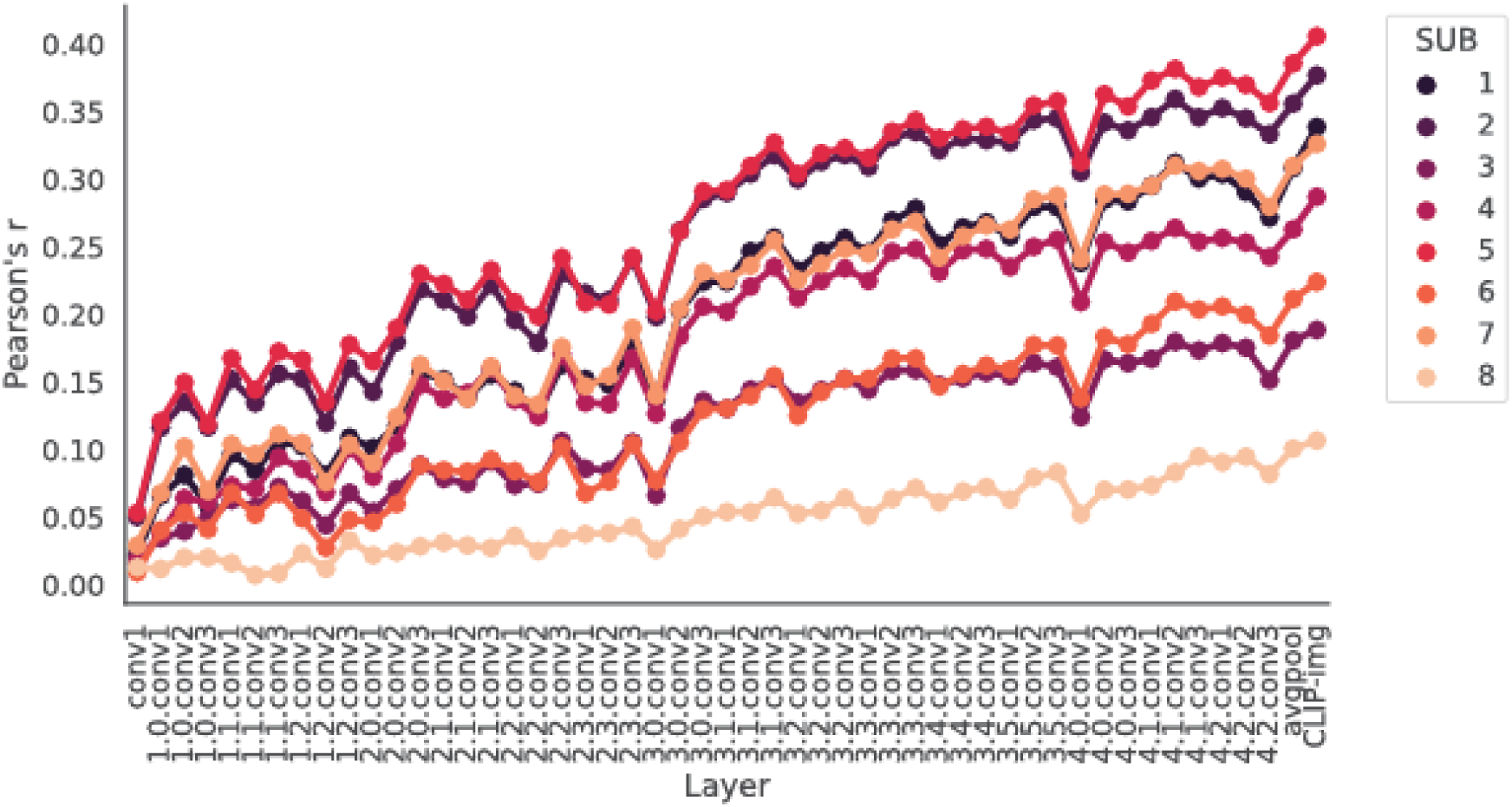
Encoding models based on features extracted from ResNet50 layers[37] also showed robust prediction of LPFC activities. We chose to report the results based on the CLIP-img features in the main text because the CLIP-img-based prediction outperforms all the ResNet50 layrs. The cross-validated prediction pipeline is the same as the CLIP-img-based pipeline described in the Methods section. The only difference between the CLIP-based and ResNet50-based pipeline is that while all 512 dimensions of CLIP-img were used, for the ResNet50 layers, we reduced the dimensions of each layer to the minimum number of components that captures more than 95 % variance in 8000 randomly sampled images from the NSD stimuli to lessen the computational demand posed by the much higher dimensionality of ResNet50 layers (ranging from 2048 to 802,816 depending on the layer).

**Figure S3:**
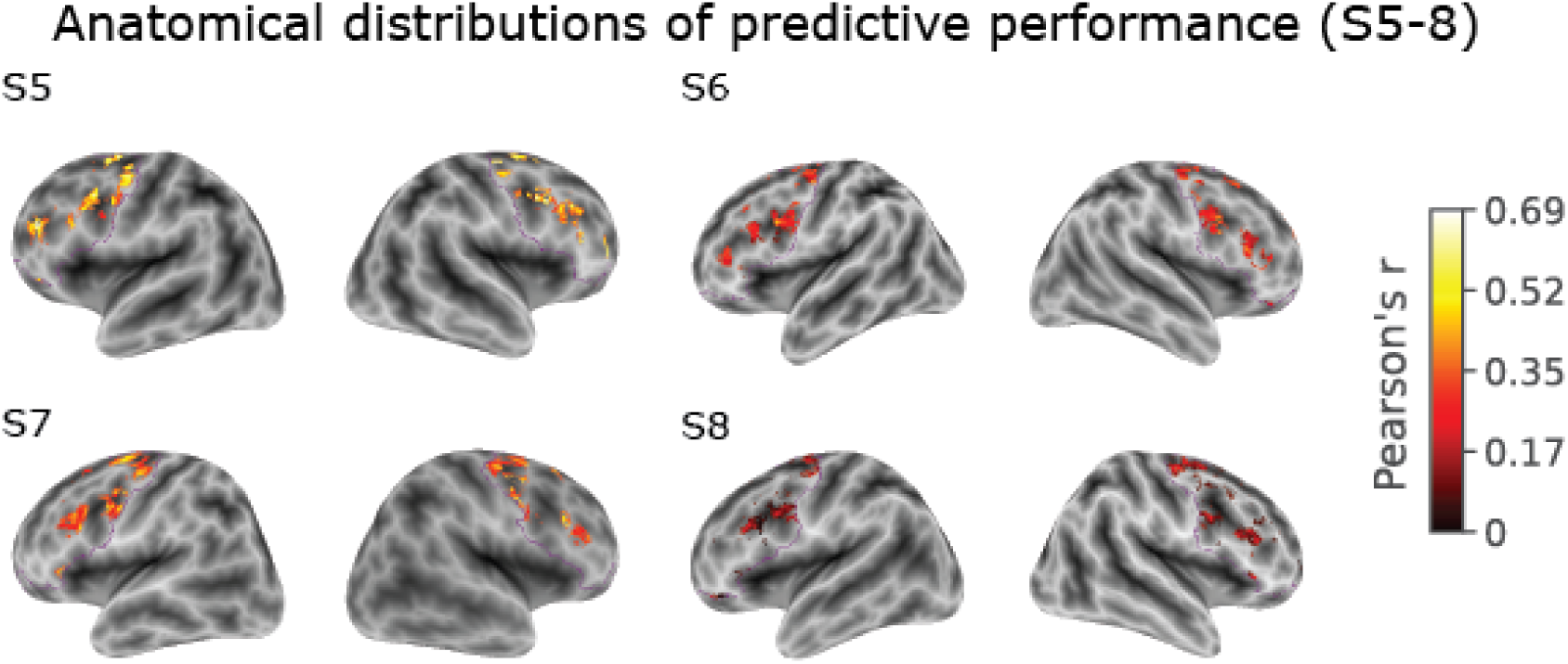
Anatomical distributions of the predictive performance based on CLIP-img features in the remaining 4 subjects. The contour of the LPFC mask is marked in purple.

**Figure S4:**
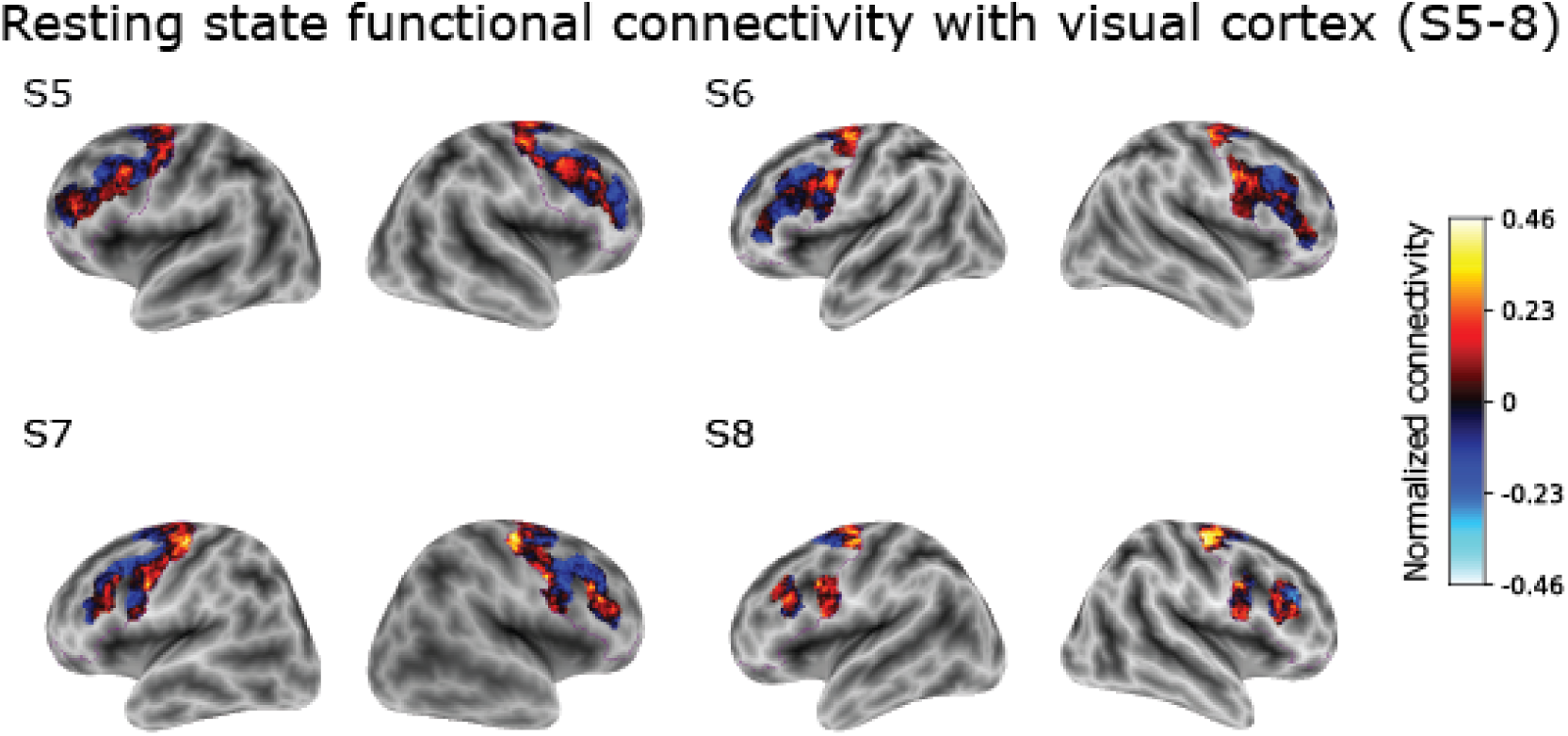
Anatomical distributions of the normalized resting state functional connectivity values between LPFC vertices and visual regions in 4 example subjects. The contour of the LPFC mask is marked in purple.

**Figure S5:**
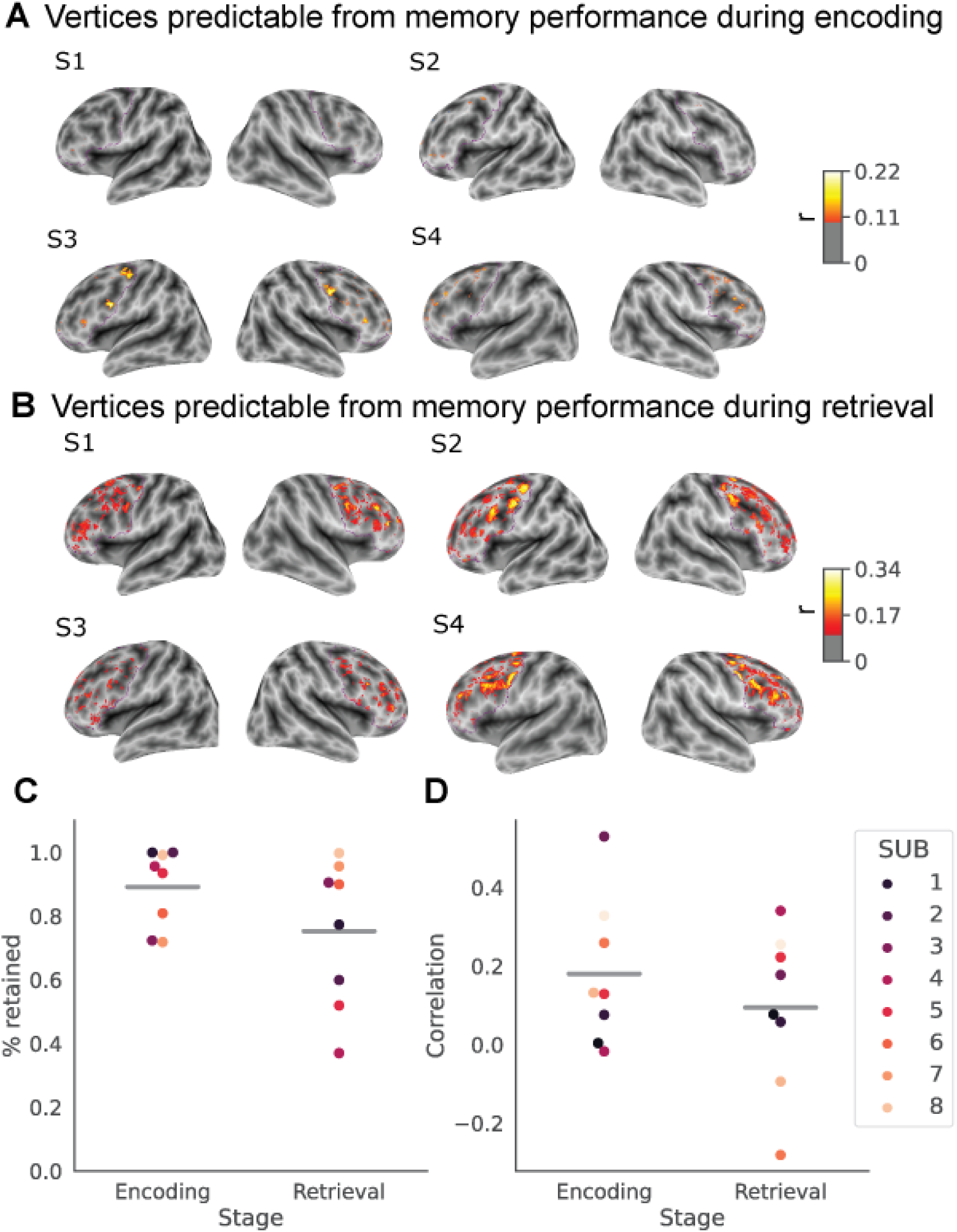
LPFC vertices that are predictable from CLIP features cannot be fully explained by memory performance. To identify brain activities associated with memory performance, we built vertex-wise encoding models using memory performance (instead of CLIP features), adopting the same cross-validation pipeline in Figure 1A. (A) Anatomical distributions of the vertices predictable from encoding performance in four example subjects (thresholded at cross-validated Pearson’s r > 0.1). For memory encoding, we focus on brain activities during the first presentation of a given image and use the memory performance on the first repetition (i.e., the second presentation) of the same image (i.e., hit or miss) and the delay between the first and second presentation as the regressors. The contour of the LPFC mask is marked in purple. (B) Anatomical distributions of the vertices predictable from retrieval performance in four example subjects (thresholded at cross-validated Pearson’s r > 0.1). For memory retrieval, we focus on brain activities during the second and/or third presentation of a given image and use the memory performance on the corresponding repetition of the same image (i.e., hit or miss) and the delay between the current and last presentation as the regressors. The contour of the LPFC mask is marked in purple. (C) Proportion of CLIP-predictable vertices retained after excluding those that also track successful memory encoding or retrieval. Each dot represents a subject. The gray line represents the mean across all subjects. Overall, the majority of vertices are retained (M*_encoding_* = 89%, M*_retrieval_* = 75%). (D) The correlations between CLIP-based predictive performance and memory-based predictive performance among CLIP-based predictable vertices (cross-validated Pearson’s r > 0.1). Each dot represents a subject. The gray line represents the mean across all subjects. Overall, the correlations are low (M*_encoding_* = 0.18, M*_retrieval_* = 0.09).

**Figure S6:**
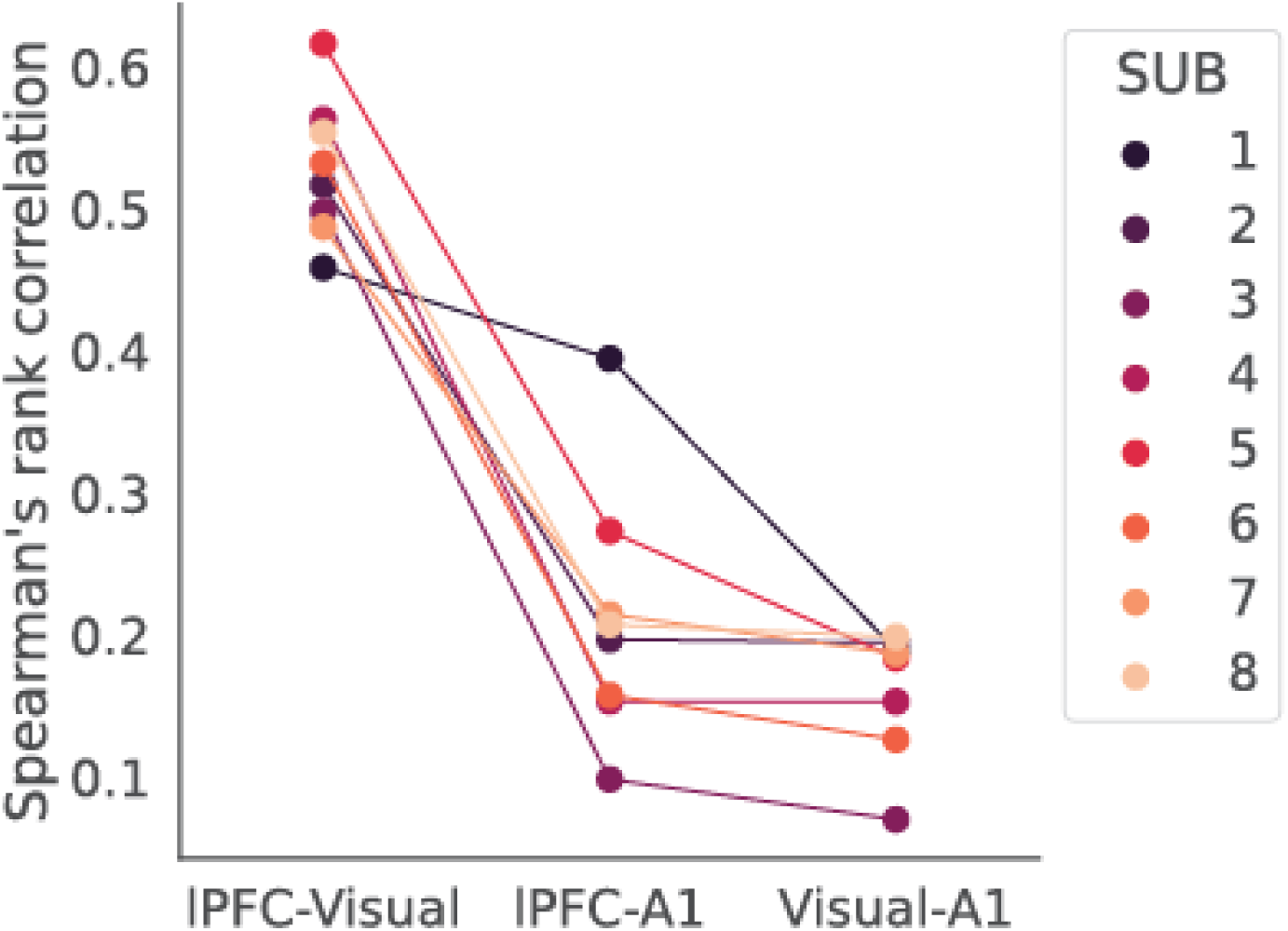
The coding schemes of LPFC and ventral visual regions are somewhat consistent within the same individuals. To demonstrate that this effect is not a global effect on any brain region pairs coming from the same individuals, we also compared the representational structures in a control region, A1 (the primary auditory cortex), where we should not expect strong visual coding, to those in lPFC and ventral visual regions respectively. The lPFC-Visual consistency is significantly higher than the lPFC-A1 and Visual-A1 consistency (both *p*s < .01 based on Wilxoncon signed rank tests), demonstrating that the observed degree of consistency between the representational structures in LPFC and ventral visual stream reflects some preservation of coding schemes for visual stimuli within the same individuals.

**Figure S7:**
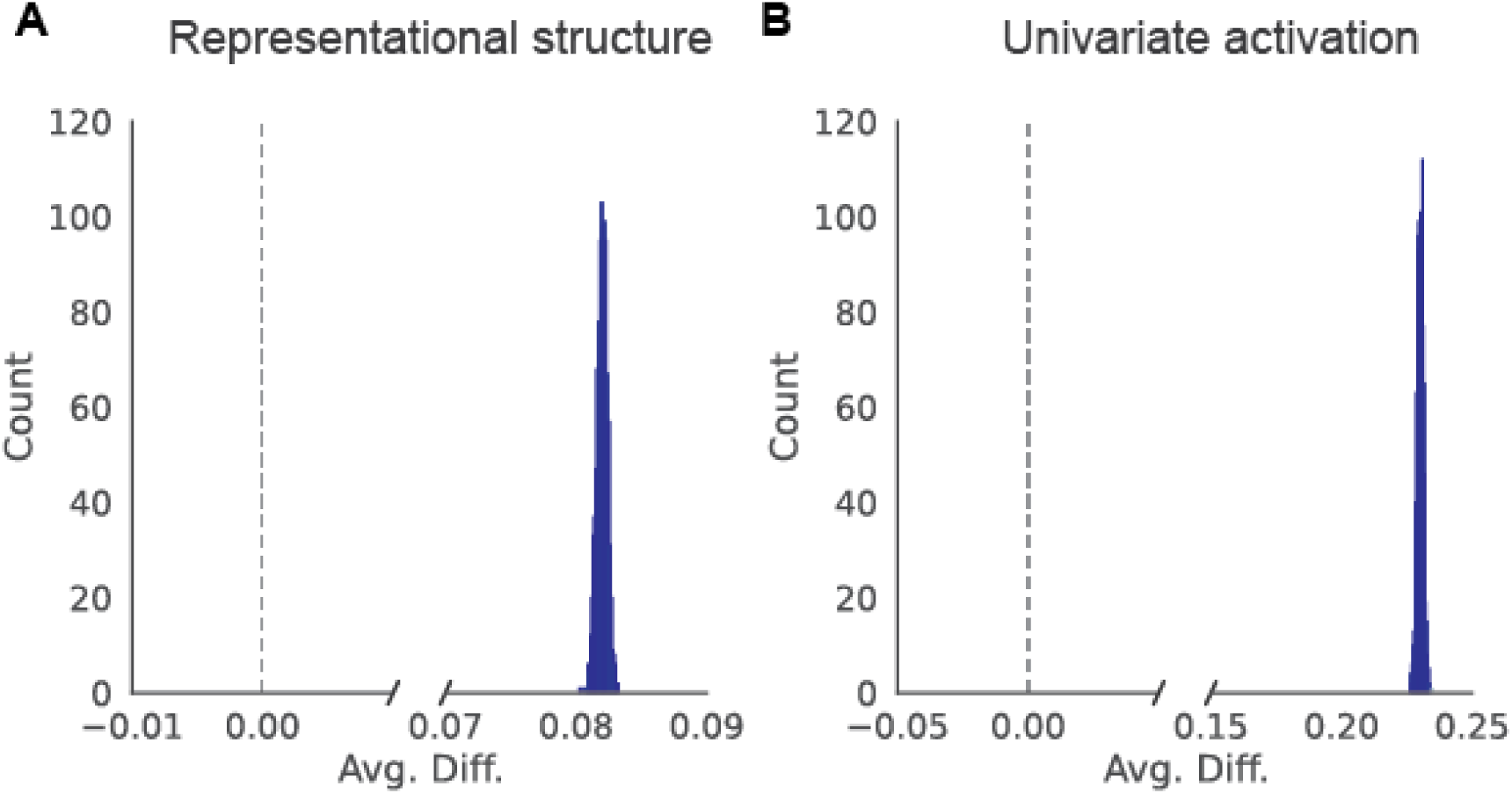
The coding scheme of LPFC is more variable across individuals compared to ventral visual regions even after we matched the overall predictability and number of vertices between the two ROIs. To ensure that the amplified individual differences in LPFC coding is not simply a result of the higher predictability and larger size of the ventral visual regions than LPFC, we subsampled vertices in ventral visual regions to match the number of included vertices as well as the range and mean of cross-validated Pearson’s r values in LPFC (i.e., mean difference in cross-validated Pearson’s r between the subsampled visual vertices and LPFC < 0.06 within each subject). We then recalculated the across-subject similarity of representational structures and predicted activation following procedures shown in Figures. 3A and 4A, respectively. We repeated this subsampling procedure 1000 times. (A) Histogram showing the distribution of the average difference between the across-subject similarity of representational similarity matrices derived based on LPFC vertices and that based on the ventral visual regions across 1000 subsampling iterations. (B) Histogram showing the distribution of the average difference between the across-subject similarity of predicted average LPFC activation and that of the predicted ventral visual regions activation across 1000 subsampling iterations.

**Figure S8:**
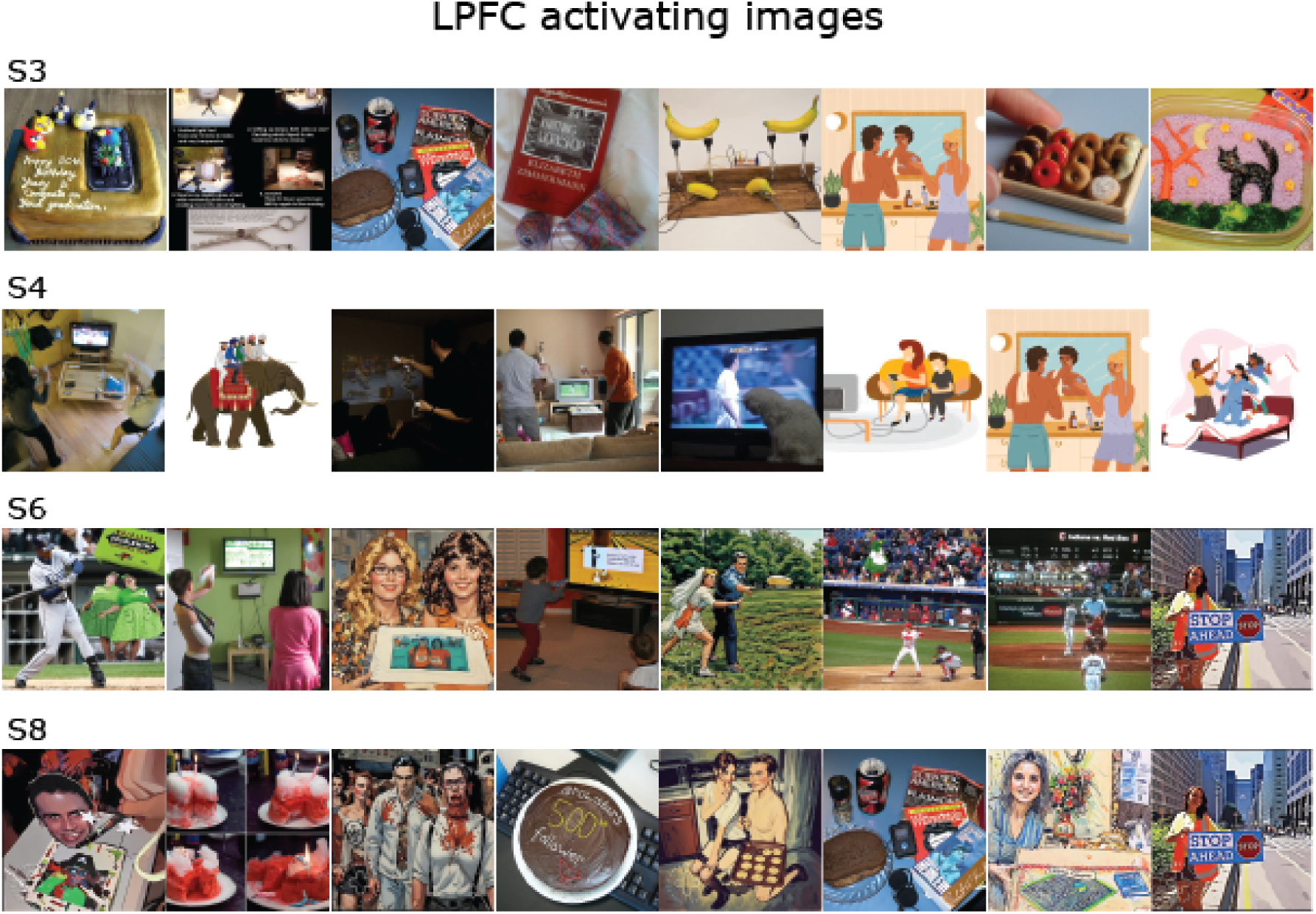
The 8 images with the largest predicted LPFC responses in 4 remaining subjects. Images with recognizable human faces have been replaced by cartoons of similar content or cartoonized with befunky (befunky.com) to comply with the policy of the bioRxiv server.

**Figure S9:**
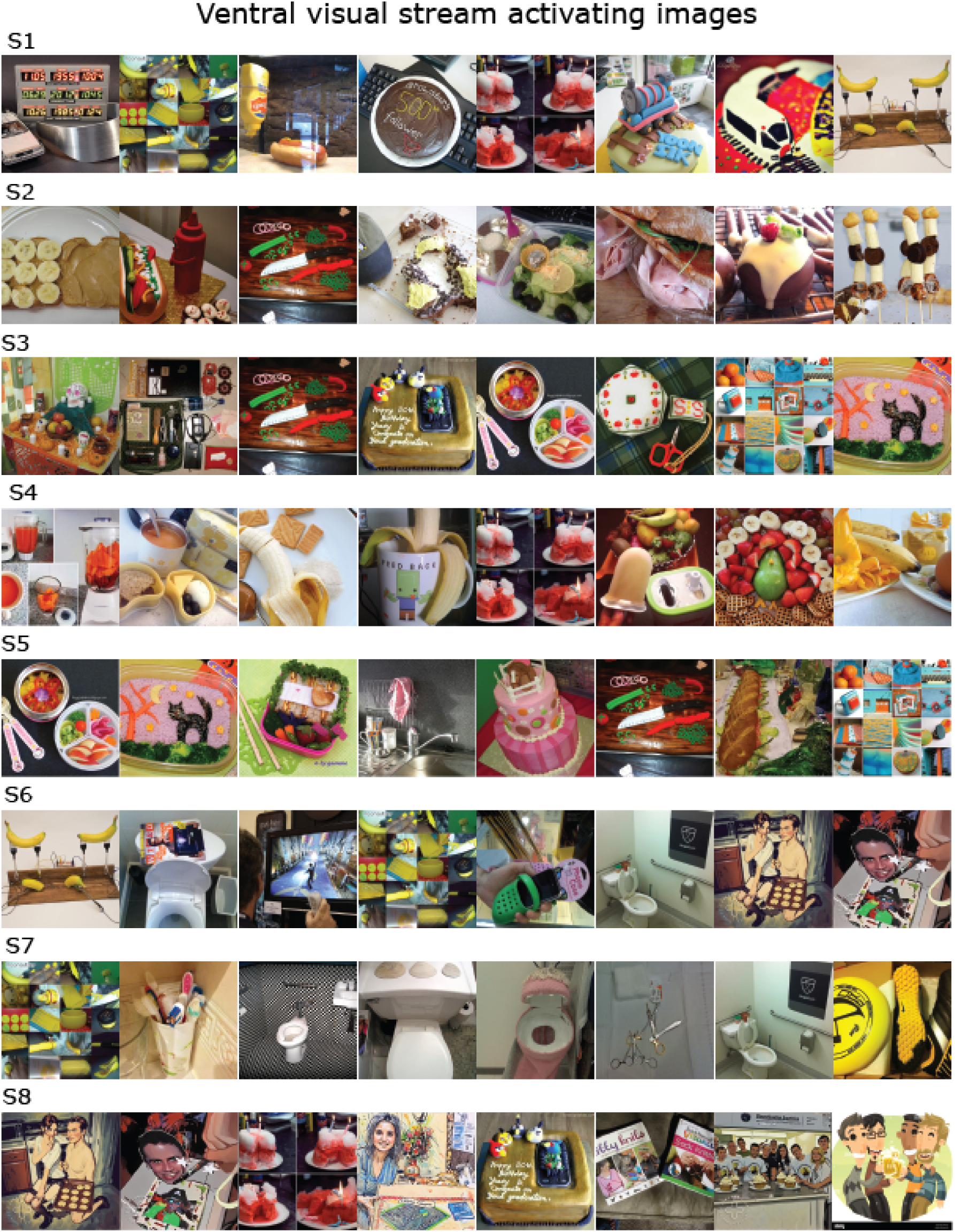
The 8 images with the largest predicted responses in ventral visual regions in all 8 subjects. Images with recognizable human faces have been replaced by cartoons of similar content or cartoonized with befunky (befunky.com) to comply with the policy of the bioRxiv server.

**Figure S10:**
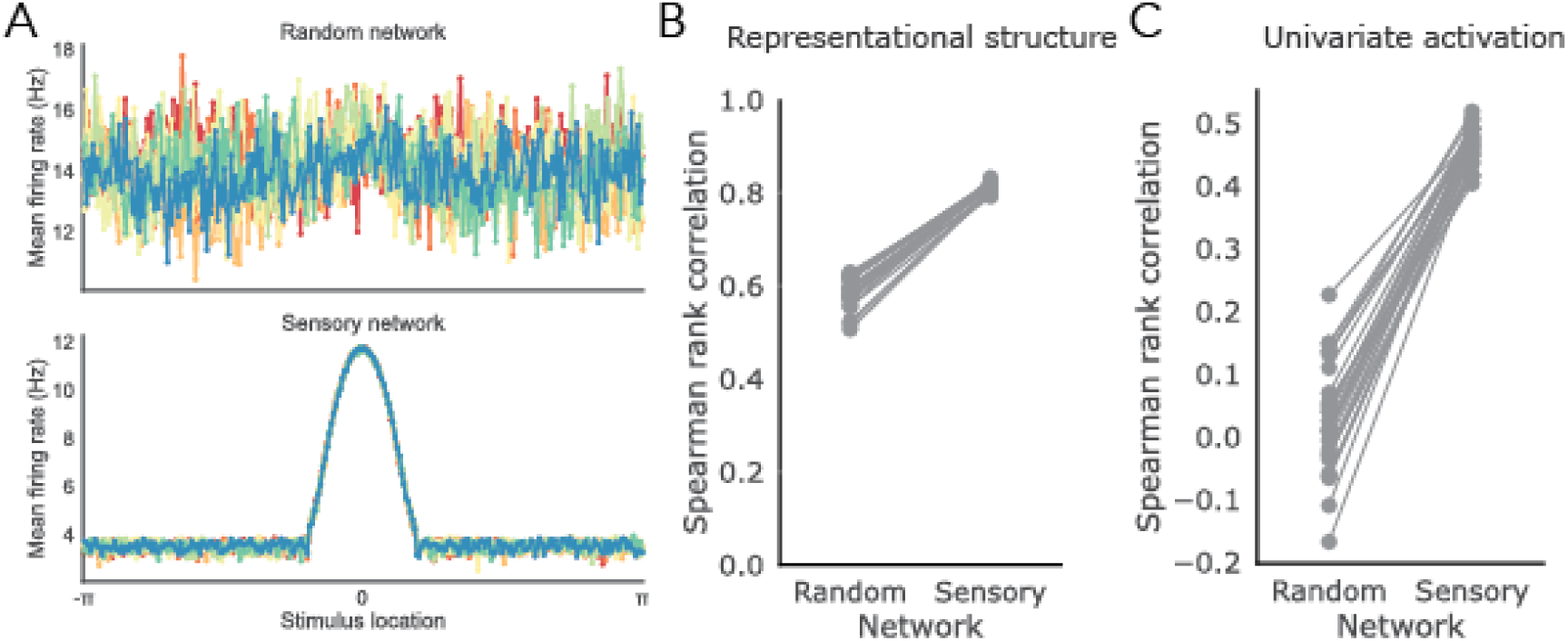
Simulation results (with one preferred stimulus location and 8 different simulated subjects) from a setup where the connections between sensory and random layers are fully random. We observed qualitatively similar results to the setup with minimal structure, both in terms of representational structures(B) and univariate activation(C). However, as observed in averaged firing rates to different stimuli in random(top) and sensory(bottom) layers shown in (A), although the sensory network shows higher overall activity to the preferred location, no preference is observed in the random network activity and thus the consistency of the univariate activation profiles in the random layer across simulated ‘subjects’ is close to zero (see C).

**Figure S11:**
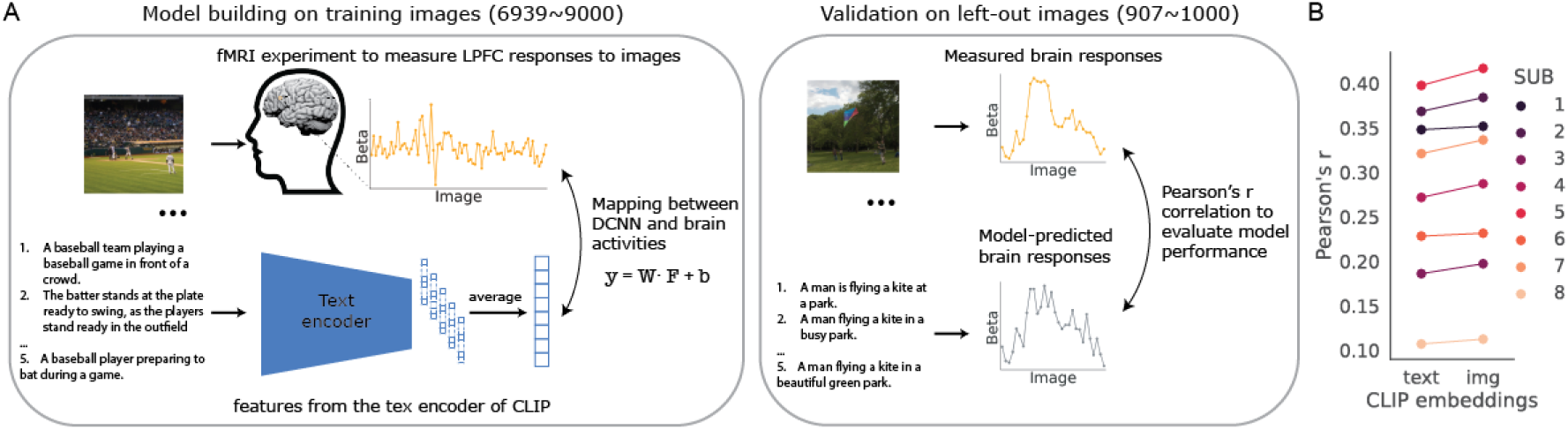
Encoding models based on the CLIP-text features of human-generated captions corresponding to the images also capture substantial variance in LPFC activities, although we still see an improvement in prediction performance from using the CLIP-img features in every single subject. (A) Schematic of the cross-validated prediction pipeline based on CLIP-text features. The only difference between the CLIP-text-based pipeline and the CLIP-img-based pipeline shown in Figure 1A is that the encoder. Instead of extracting CLIP-img embeddings using images as input, we extract the CLIP-text embeddings using the captions corresponding to the presented images provided by the COCO dataset[74]. Note that because each image comes with 5 captions from different human observers, we simply average the text embeddings corresponding to the same image to generate a mean text embedding for each image. (B) Average Pearson’s r between predicted and observed responses in the top 10% LPFC vertices in the held-out data, using CLIP-text or CLIP-img embeddings. Each dot represents a subject.

## References

1. Stanovich KE and West RF. Individual differences in rational thought. Journal of experimental psychology: general 1998;127:161.

2. Gray ME and Holyoak KJ. Individual differences in relational reasoning. Memory & cognition 2020;48:96–110.

3. Schulz L, Rollwage M, Dolan RJ, and Fleming SM. Dogmatism manifests in lowered information search under uncertainty. Proceedings of the National Academy of Sciences 2020;117:31527–34.

4. Preuss TM and Wise SP. Evolution of prefrontal cortex. Neuropsychopharmacology 2022;47:3– 19.

5. Mueller S, Wang D, Fox MD, et al. Individual variability in functional connectivity architecture of the human brain. Neuron 2013;77:586–95.

6. Finn ES, Shen X, Scheinost D, et al. Functional connectome fingerprinting: identifying individuals using patterns of brain connectivity. Nature neuroscience 2015;18:1664–71.

7. Cole MW, Yarkoni T, Repov̌s G, Anticevic A, and Braver TS. Global connectivity of prefrontal cortex predicts cognitive control and intelligence. Journal of Neuroscience 2012;32:8988–99.

8. Fuster J. The prefrontal cortex. Academic press, 2015.

9. Duncan J. The multiple-demand (MD) system of the primate brain: mental programs for intelligent behaviour. Trends in cognitive sciences 2010;14:172–9.

10. Miller EK and Cohen JD. An integrative theory of prefrontal cortex function. Annual review of neuroscience 2001;24:167–202.

11. Passingham RE and Wise SP. The neurobiology of the prefrontal cortex: anatomy, evolution, and the origin of insight. OUP Oxford, 2012.

12. Fedorenko E and Varley R. Language and thought are not the same thing: evidence from neuroimaging and neurological patients. Annals of the New York Academy of Sciences 2016;1369:132– 53.

13. Rigotti M, Barak O, Warden MR, et al. The importance of mixed selectivity in complex cognitive tasks. Nature 2013;497:585–90.

14. Fusi S, Miller EK, and Rigotti M. Why neurons mix: high dimensionality for higher cognition. Current opinion in neurobiology 2016;37:66–74.

15. Tye KM, Miller EK, Taschbach FH, Benna MK, Rigotti M, and Fusi S. Mixed selectivity: Cellular computations for complexity. Neuron 2024.

16. Hubel DH and Wiesel TN. Receptive fields, binocular interaction and functional architecture in the cat’s visual cortex. The Journal of physiology 1962;160:106.

17. Kornblith S and Tsao DY. How thoughts arise from sights: inferotemporal and prefrontal contributions to vision. Current Opinion in Neurobiology 2017;46:208–18.

18. Riley MR, Qi XL, and Constantinidis C. Functional specialization of areas along the anterior– posterior axis of the primate prefrontal cortex. Cerebral Cortex 2017;27:3683–97.

19. Tsao DY, Schweers N, Moeller S, and Freiwald WA. Patches of face-selective cortex in the macaque frontal lobe. Nature neuroscience 2008;11:877–9.

20. Haile TM, Bohon KS, Romero MC, and Conway BR. Visual stimulus-driven functional organization of macaque prefrontal cortex. Neuroimage 2019;188:427–44.

21. Rose O and Ponce CR. A concentration of visual cortex-like neurons in prefrontal cortex. Nature Communications 2024;15:7002.

22. Freedman DJ, Riesenhuber M, Poggio T, and Miller EK. A comparison of primate prefrontal and inferior temporal cortices during visual categorization. Journal of Neuroscience 2003;23:5235–46.

23. Fyall AM, El-Shamayleh Y, Choi H, Shea-Brown E, and Pasupathy A. Dynamic representation of partially occluded objects in primate prefrontal and visual cortex. Elife 2017;6:e25784.

24. Kar K and DiCarlo JJ. Fast recurrent processing via ventrolateral prefrontal cortex is needed by the primate ventral stream for robust core visual object recognition. Neuron 2021;109:164–76.

25. Brascamp J, Sterzer P, Blake R, and Knapen T. Multistable perception and the role of the frontoparietal cortex in perceptual inference. Annual review of psychology 2018;69:77–103.

26. Wang M, Arteaga D, and He BJ. Brain mechanisms for simple perception and bistable perception. Proceedings of the National Academy of Sciences 2013;110:E3350–E3359.

27. Ge Y, Taschereau-Dumouchel V, Lin Q, Moharramipour A, Sun Z, and Lau H. Representation of visual uniformity in the lateral prefrontal cortex. bioRxiv 2024:2024–7.

28. Watanabe T. Causal roles of prefrontal cortex during spontaneous perceptual switching are determined by brain state dynamics. Elife 2021;10:e69079.

29. Wang AY, Kay K, Naselaris T, Tarr MJ, and Wehbe L. Better models of human high-level visual cortex emerge from natural language supervision with a large and diverse dataset. Nature Machine Intelligence 2023;5:1415–26.

30. Huth AG, Nishimoto S, Vu AT, and Gallant JL. A continuous semantic space describes the representation of thousands of object and action categories across the human brain. Neuron 2012;76:1210–24.

31. Khosla M, Ngo GH, Jamison K, Kuceyeski A, and Sabuncu MR. Cortical response to naturalistic stimuli is largely predictable with deep neural networks. Science Advances 2021;7:eabe7547.

32. Allen EJ, St-Yves G, Wu Y, et al. A massive 7T fMRI dataset to bridge cognitive neuroscience and artificial intelligence. Nature neuroscience 2022;25:116–26.

33. Yamins DL and DiCarlo JJ. Using goal-driven deep learning models to understand sensory cortex. Nature neuroscience 2016;19:356–65.

34. Lindsay GW. Convolutional neural networks as a model of the visual system: Past, present, and future. Journal of cognitive neuroscience 2021;33:2017–31.

35. Golan T, Taylor J, Schütt H, et al. Deep neural networks are not a single hypothesis but a language for expressing computational hypotheses. Behavioral and Brain Sciences 2023;46.

36. Radford A, Kim JW, Hallacy C, et al. Learning transferable visual models from natural language supervision. In: International conference on machine learning. PMLR. 2021:8748–63.

37. He K, Zhang X, Ren S, and Sun J. Deep residual learning for image recognition. In: Proceedings of the IEEE conference on computer vision and pattern recognition. 2016:770–8.

38. Glasser MF, Coalson TS, Robinson EC, et al. A multi-modal parcellation of human cerebral cortex. Nature 2016;536:171–8.

39. Ratan Murty NA, Bashivan P, Abate A, DiCarlo JJ, and Kanwisher N. Computational models of category-selective brain regions enable high-throughput tests of selectivity. Nature communications 2021;12:5540.

40. Bouchacourt F and Buschman TJ. A flexible model of working memory. Neuron 2019;103:147–60.

41. Kanwisher N, McDermott J, and Chun MM. The Fusiform Face Area: A Module in Human Extrastriate Cortex Specialized for Face Perception. The Journal of Neuroscience 1997;17:4302–11.

42. Koch C, Massimini M, Boly M, and Tononi G. Neural correlates of consciousness: progress and problems. Nature reviews neuroscience 2016;17:307–21.

43. Yeterian EH, Pandya DN, Tomaiuolo F, and Petrides M. The cortical connectivity of the prefrontal cortex in the monkey brain. Cortex 2012;48:58–81.

44. Fleming SM, Huijgen J, and Dolan RJ. Prefrontal Contributions to Metacognition in Perceptual Decision Making. The Journal of Neuroscience 2012;32:6117–25.

45. McCurdy LY, Maniscalco B, Metcalfe J, Liu KY, Lange FP de, and Lau H. Anatomical Coupling between Distinct Metacognitive Systems for Memory and Visual Perception. The Journal of Neuroscience 2013;33:1897–906.

46. Morales J, Lau H, and Fleming SM. Domain-General and Domain-Specific Patterns of Activity Supporting Metacognition in Human Prefrontal Cortex. The Journal of Neuroscience 2018;38:3534–46.

47. Middlebrooks PG and Sommer MA. Neuronal Correlates of Metacognition in Primate Frontal Cortex. Neuron 2012;75:517–30.

48. Berke M, Azerbayev Z, Belledonne M, Tavares Z, and Jara-Ettinger J. MetaCOG: A Heirar-chical Probabilistic Model for Learning Meta-Cognitive Visual Representations. In: The 40th Conference on Uncertainty in Artificial Intelligence. 2024.

49. Himmelberg MM, Winawer J, and Carrasco M. Linking individual differences in human primary visual cortex to contrast sensitivity around the visual field. Nature Communications 2022;13.

50. Moutsiana C, Haas B de, Papageorgiou A, et al. Cortical idiosyncrasies predict the perception of object size. Nature Communications 2016;7.

51. Charest I, Kievit RA, Schmitz TW, Deca D, and Kriegeskorte N. Unique semantic space in the brain of each beholder predicts perceived similarity. Proceedings of the National Academy of Sciences 2014;111:14565–70.

52. Lee J and Geng JJ. Idiosyncratic Patterns of Representational Similarity in Prefrontal Cortex Predict Attentional Performance. The Journal of Neuroscience 2016;37:1257–68.

53. Wang G, Foxwell MJ, Cichy RM, Pitcher D, and Kaiser D. Individual differences in internal models explain idiosyncrasies in scene perception. Cognition 2024;245:105723.

54. Fecteau J and Mimpz D. Salience, relevance, and firing: a priority map for target selection. Trends in Cognitive Sciences 2006;10:382–90.

55. Awh E, Belopolsky AV, and Theeuwes J. Top-down versus bottom-up attentional control: a failed theoretical dichotomy. Trends in Cognitive Sciences 2012;16:437–43.

56. Arnatkeviciute A, Fulcher BD, Oldham S, et al. Genetic influences on hub connectivity of the human connectome. Nature Communications 2021;12:4237.

57. Thompson PM, Cannon TD, Narr KL, et al. Genetic influences on brain structure. Nature neuroscience 2001;4:1253–8.

58. Balas B and Saville A. Hometown size affects the processing of naturalistic face variability. Vision research 2017;141:228–36.

59. Coutrot A, Manley E, Goodroe S, et al. Entropy of city street networks linked to future spatial navigation ability. Nature 2022;604:104–10.

60. Sugihara T, Diltz MD, Averbeck BB, and Romanski LM. Integration of auditory and visual communication information in the primate ventrolateral prefrontal cortex. Journal of Neuroscience 2006;26:11138–47.

61. Watanabe M. Frontal units of the monkey coding the associative significance of visual and auditory stimuli. Experimental brain research 1992;89:233–47.

62. Doerig A, Kietzmann TC, Allen E, et al. Visual representations in the human brain are aligned with large language models. arXiv preprint arXiv:2209.11737 2022.

63. Shoham A, Broday-Dvir R, Yaron I, Yovel G, and Malach R. Text-related functionality of visual human pre-frontal activations revealed through neural network convergence. bioRxiv 2024:2024–4.

64. Haxby JV, Guntupalli JS, Nastase SA, and Feilong M. Hyperalignment: Modeling shared information encoded in idiosyncratic cortical topographies. elife 2020;9:e56601.

65. Chen PHC, Chen J, Yeshurun Y, Hasson U, Haxby J, and Ramadge PJ. A reduced-dimension fMRI shared response model. Advances in neural information processing systems 2015;28.

66. Poldrack RA. Precision neuroscience: dense sampling of individual brains. Neuron 2017;95:727–9.

67. Kupers ER, Knapen T, Merriam EP, and Kay KN. Principles of intensive human neuroimaging. Trends in Neurosciences 2024.

68. Lee HJ, Dworetsky A, Labora N, and Gratton C. Using precision approaches to improve brain-behavior prediction. Trends in Cognitive Sciences 2025;29:170–83.

69. Wang L, Mruczek RE, Arcaro MJ, and Kastner S. Probabilistic maps of visual topography in human cortex. Cerebral cortex 2015;25:3911–31.

70. Murray LJ and Ranganath C. The dorsolateral prefrontal cortex contributes to successful relational memory encoding. Journal of Neuroscience 2007;27:5515–22.

71. Badre D and Wagner AD. Left ventrolateral prefrontal cortex and the cognitive control of memory. Neuropsychologia 2007;45:2883–901.

72. Long NM, Öztekin I, and Badre D. Separable prefrontal cortex contributions to free recall. Journal of Neuroscience 2010;30:10967–76.

73. Paller KA and Wagner AD. Observing the transformation of experience into memory. Trends in cognitive sciences 2002;6:93–102.

74. Lin TY, Maire M, Belongie S, et al. Microsoft COCO: Common Objects in Context. In: Computer Vision – ECCV 2014. Springer International Publishing, 2014:740–55. doi: 10.1007/978-3-319-10602-1_48. url: http://dx.doi.org/10.1007/978-3-319-10602-1_48.

